# Unravelling bacteriochlorophyll biosynthesis in the uncultured *Roseilinea*

**DOI:** 10.64898/2026.11.16.766355

**Authors:** Nicola M Rudling, Dowrung Namoon, Vera Thiel, Daniel P Canniffe

## Abstract

Filamentous anoxygenic phototrophs (FAPs) of the *Chloroflexota* phylum are abundant members of microbial mat communities. All isolated and culturable FAPs are found in a single class within this phylum, but metagenomic studies of hot spring communities from around the world have indicated that phototrophic members of an additional class may exist, which have so far eluded cultivation for study in the laboratory. Metagenome-assemble genomes (MAGs) of these “*Candidatus* Roseilinea” species all lack genes encoding the central enzymes of bacteriochlorophyll biosynthesis common to all FAPs and to the seven other diverse phyla of anoxygenic phototrophs, which could indicate that these organisms are no longer phototrophic or that they employ a novel set of enzymes to generate chlorophyllide from magnesium protoporphyrin IX. Here we demonstrate that the five terminal enzymes of bacteriochlorophyll biosynthesis encoded in the MAG of Japanese hot spring-dwelling “*Ca*. Roseilinea sp. NK_OTU-006” are active and restore pigment production and phototrophic growth in a set of mutants of the model purple phototrophic bacterium *Rhodobacter sphaeroides*. While many organisms are known to require exogenous supply of metabolically costly modified tetrapyrroles such as vitamin B_12_ for growth, the retained function of the tested *Roseilinea* enzymes and the restriction of these species to dense microbial mats leads us to hypothesise that these organisms may be the first examples of bacteriochlorophyll auxotrophs. We suggest that *Roseilinea* scavenge pigment precursors and/or breakdown products excreted by other members of the mat to generate the mature bacteriochlorophyll they require to perform phototrophy, an exchange that we demonstrate can occur between *Rhodobacter sphaeroides* mutants. We further suggest that future attempts to isolate *Roseilinea* species should employ the exogenous supplementation of (bacterio)chlorophyllide into culture medium.

## INTRODUCTION

Chlorophototrophic organisms use chlorophyll (Chl) and/or bacteriochlorophyll (BChl) pigments to capture solar radiation and convert it to chemical energy to power cellular metabolism (Bryant and Canniffe 2018). The biosynthesis of these pigments involves the reduction of the C17=C18 double bond of the porphyrin precursor 8-vinyl (8V) protochlorophyllide (PChlide) to produce 8V chlorophyllide (Chlide), a chlorin that is the universal precursor for all known, naturally-occurring (B)Chl pigments (Bryant et al. 2020) (**Fig. 1**). This reaction is catalysed by two unrelated PChlide oxidoreductase (POR) enzymes, each employing different chemistries. Chl-containing oxygenic phototrophs primarily use a single-subunit, light-activated enzyme, LPOR, which uses NADPH as the reductant (Heyes and Hunter 2005); angiosperms rely solely on this enzyme to synthesise Chl, and its inactivity in the dark halts biosynthesis in flowering plants (Apel et al. 1998). The majority of BChl-containing anoxygenic phototrophs utilise a multi-subunit enzyme that can operate in the dark, DPOR, that derives electrons from ferredoxin at the BchL protein, and shuttles them to the substrate-binding BchN–BchB complex via multiple [4Fe–4S] cluster cofactors, an enzyme that is thought to have evolved from nitrogenase (Muraki et al. 2010). Oxygenic cyanobacteria, algae and gymnosperms have both enzymes, using DPOR to maintain Chl levels at night (Fujita and Bauer 2003). Some aerobic anoxygenic phototrophs—obligately aerobic bacteria that supplement their metabolism with anoxygenic photosynthesis—commonly use LPOR acquired from a cyanobacterium, presumably because the oxygen-labile [4Fe–4S] cofactors in DPOR cannot function in the host’s oxic environment (Kaschner et al. 2014; Kasalický et al. 2018).

**Figure 1.**
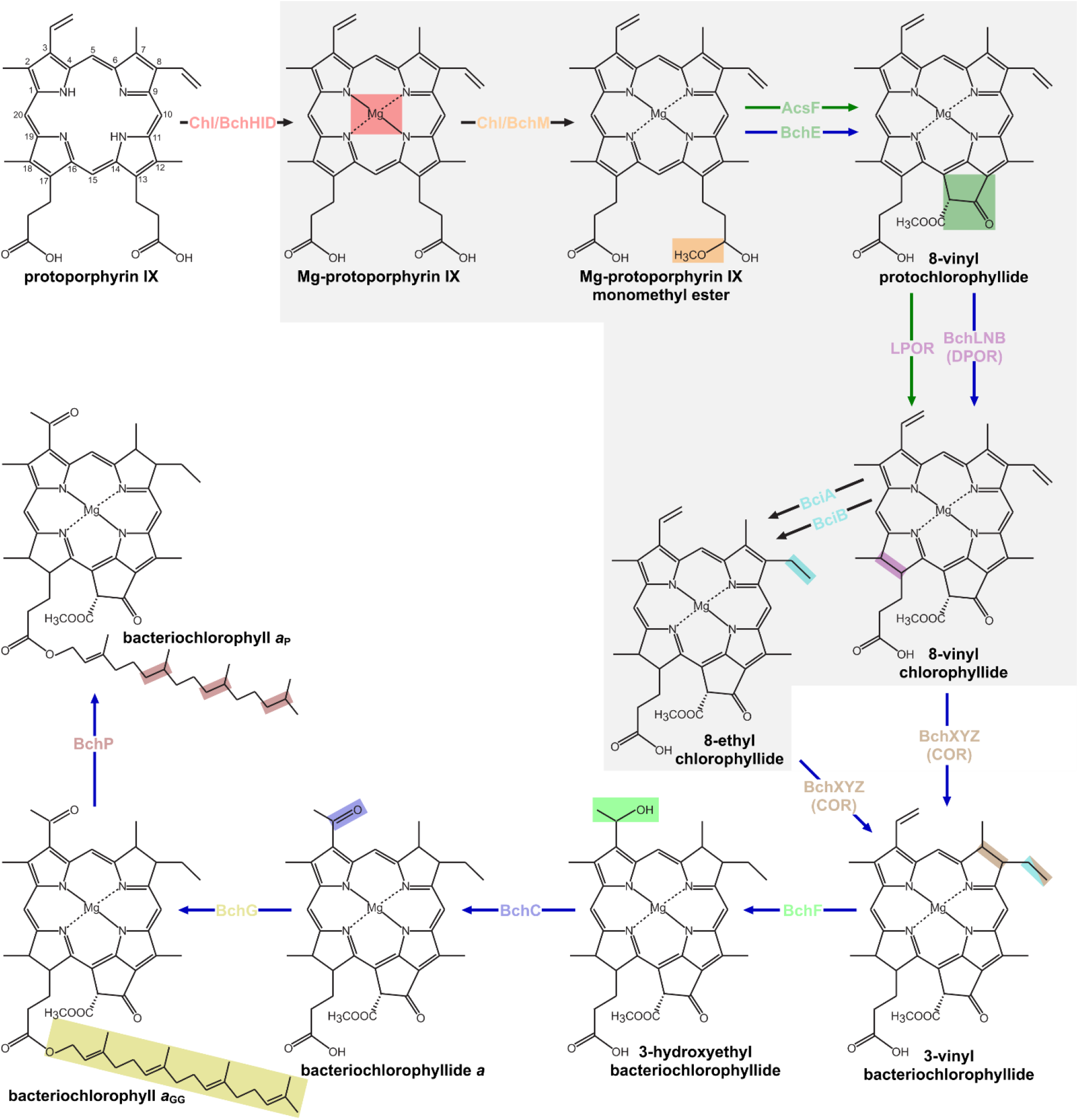
Biosynthesis of precursors of Chl *a* and BChl *a*. IUPAC numbering of macrocycle carbons is indicated on protoporphyrin IX. Major enzymes/enzyme subunits are overlaid on reaction arrows, and catalysed modifications are highlighted on molecular structures. Arrows indicating primary routes for Chl or BChl biosynthesis are coloured green or blue, respectively. Black arrows indicate modification by enzyme common to both pathways. Dashed arrows indicate multiple biosynthetic steps. Stimulatory/minor enzyme subunits are omitted for clarity. Note that COR is able to catalyse reduction at C8-vinyl, as well as C7=C8. The biosynthetic steps apparently missing in phototrophic *Roseilinea* are on grey background.

The first step extending from biosynthesis of Chls to the true BChls is the reduction of the C7=C8 double bond of a chlorophyllide (Chlide) pigment to a bacteriochlorophyllide (BChlide), converting the chlorin to a bacteriochlorin (see **Fig. 1**). This reaction is catalysed by Chlide oxidoreductase (COR), made up of BchX and BchY–BchZ components, each sharing high sequence identity with the BchL, BchN, and BchB subunits of DPOR respectively, and using similar chemistry (Burke et al. 1993a; Kiesel et al. 2015). It is likely that a gene duplication event resulted in the evolution of these two paralogous enzymes (Burke et al. 1993b). Two isoforms of COR are known to exist; the enzyme found in organisms that make BChl *a* (COR*a*) has an additional C8-vinyl reductase activity, allowing it to produce a pigment with a C8 ethyl group using Chlide substrates carrying C8 vinyl or ethyl substituents, whereas the enzyme in organisms making BChl *b* or BChl *g* (COR*b*) lack this activity and only use C8-vinyl Chlide as a substrate (Tsukatani et al. 2013; Canniffe and Hunter 2014). The vast majority of BChl *a*-containing organisms studied to date contain an additional, canonical C8-vinyl reductase, encoded by either *bciA* or *bciB*, which may act as a failsafe to ensure ethyl-carrying BChl *a* is formed in these cells.

Anoxygenic phototrophs are found in diverse bacterial phyla throughout the tree of life (Hug 2024; White et al. 2025). Phototrophic members of the *Pseudomonadota* (proteobacteria known commonly as purple bacteria), *Bacillota* (*Firmicutes*), *Chlorobiota* (green sulfur bacteria), and *Chloroflexota* (filamentous anoxygenic phototrophs (FAP)) have been widely studied for decades (Thweatt et al. 2019). Since the turn of the century, anoxygenic phototrophic members of the *Acidobacteriota*, *Gemmatimonadota*, and *Candidatus* (*Ca*.) Eremiobacterota have also been isolated and studied (Bryant et al. 2007; Zeng et al. 2014; Yabe et al. 2022), and genes involved in anoxygenic photosynthesis have been found in metagenome assembled genomes (MAGs) of members of the *Bacteroidota* and *Myxococcota* phyla (Tahon and Willems 2017; Li et al. 2023). These extant anoxygenic phototrophs from diverse phyla use single, BChl-based photosystems comprised of few subunits to generate chemical energy, rather than the two Chl-containing, multi-component photosystems found in cyanobacteria, the only prokaryotes capable of oxygenic phototrophy, an ancestor of which being the progenitor of the chloroplast (Blankenship 2022). For these reasons, researchers have suggested that BChl-based phototrophy may have pre-dated the Chl-dependent process, despite anoxygenic phototrophs using pigments with greater structural complexity (Blankenship 1992; Xiong et al. 2000). These studies counter the classical ‘Granick hypothesis’, which postulates that biochemical pathways become progressively more complex in a stepwise fashion, and thus that Chl preceded BChl (Granick 1957, 1965). Those proposing the ‘BChl first’ hypothesis have suggested that bacteriochlorins could have arisen directly from porphyrins (e.g. BChlide from PChlide) through the activity of a promiscuous oxidoreductase that reduced both C17=C18 and C7=C8 bonds, and that gene duplication resulted in enzymes that evolved specificity (DPOR and COR) rather than the evolution of COR from POR to extend the biosynthetic pathway of these pigments (Burke et al. 1993b; Lockhart et al. 1996; Hohmann-Marriott and Blankenship 2011). Contemporary phylogenetic studies making use of the enormous genomic/metagenomic datasets now available are reverting to support the theory that BChl biosynthesis is a more recent evolutionary innovation than Chl-based phototrophy (Cardona 2016; Ward and Shih 2021). It is thought that the ancient Chl-supported process was originally anoxygenic, and that BChl biosynthesis evolved to permit phototrophy deeper in microbial mats and water columns, using far-red/near-infrared wavelengths that are not absorbed by the Chl-containing organisms at the surface (Martin et al. 2018).

All culturable FAPs isolated to date are members of the *Chloroflexia* class of *Chloroflexota*, the most widely studied being species of *Chloroflexus* and *Roseiflexus*. Metagenomic studies of hot-spring microbial mats indicate the existence of additional phototrophic members outside the *Chloroflexia* class (Klatt et al. 2011, 2013a; Thiel et al. 2016; Ward et al. 2018). “*Candidatus* Roseilinea gracilis” (formerly “*Ca*. Roseilinea gracile”, now renamed according to Oren (Oren 2024)) was the first of these to be identified and initially referred to as “*Anaerolinea*-like” (Klatt et al. 2011, 2013a; Thiel et al. 2016; Tank et al. 2017); *Anaerolinea* are non-phototrophic chemoheterotrophs in the *Chloroflexota* (Narihiro and Kamagata 2013). Although this bacterium has not been isolated into axenic culture, very long and fine cells displaying fluorescence emission associated with BChl *a* have been observed in enrichments and environmental samples. Through (meta)genome analyses, additional closely related organisms have been discovered in microbial mats worldwide (Ward et al. 2018, 2019, 2021; Martinez et al. 2020; Kato et al. 2022), and will be referred to as *Roseilinea* hereafter. Interestingly, all of these genomes lack *bchM*, *bchE*/*acsF*, *bchLNB*/*por*, and *bciA*/*bchB* but contain all other essential genes for BChl biosynthesis (e.g. *bchHID*, *bchXYX*; see **Fig. 1**) (Ward et al. 2018, 2021; Ward and Shih 2021) (**Fig. 2**). This suggest that phototrophic *Roseilinea* synthesise BChl via an atypical route that uses unidentified enzymes (Ward et al. 2021; Ward and Shih 2021). Intriguingly, the genome of the only aerobic phototrophic *Chlorobiota*, “*Ca*. Thermochlorobacter aerophilum” also lacks *bchM* and *bchE*/*acsF* genes but contains *bchLNB* and *bciB* (Liu et al. 2012). It is possible that unidentified enzymes shared between the *Roseilinea* and “*Ca*. T. aerophilum” replace BchM and BchE/AcsF in these organisms. Ward & Shih have recently suggested that the *bchXYZ* genes in *Roseilinea* may encode an enzyme able to reduce both C17=C18 and C7=C8 in the same manner as the ancient promiscuous reductase in the hypothetical ‘BChl first’ pathway (Ward and Shih 2021; Burke et al. 1993b).

**Figure 2.**
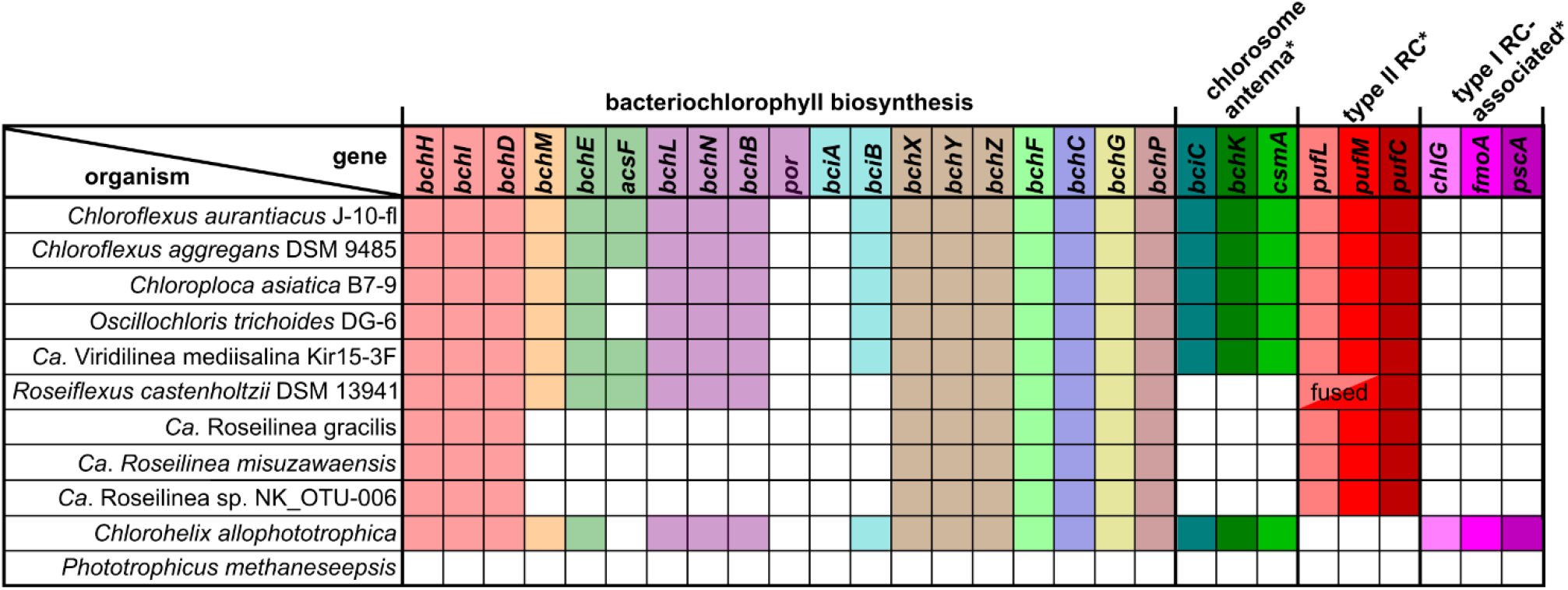
Presence of photosynthesis genes in representative FAPs. Shaded squares indicate gene is present in the genome, with bch genes coloured as in Figure 1. An asterisk (*) indicates that the genes are representative rather than exhaustive.

In the present work we test the activity of the terminal BChl biosynthesis enzymes—encoded by the *bchXYZFCGP* genes—of the recently-sequenced “*Ca*. Roseilinea sp. NK_OTU-006” in a heterologous system to reconstruct this pathway and confirm the identity of the BChl it makes, in so doing testing the possibility of it using the promiscuous PChlide/Chlide oxidoreductase, and propose a scavenging mechanism for BChl biosynthesis in these microbial mat dwelling FAPs.

## RESULTS

### Testing the activity of the *Roseilinea* COR enzyme

To investigate whether the *Roseilinea bchYZ* genes encode an enzyme with 8-vinyl reductase activity typical of COR*a*, the native *bciA* gene encoding the predominant 8-vinyl reductase was deleted from *Rba. sphaeroides* using the pK18*mobsacB* counter selection system. The deletion was confirmed by PCR performed on extracted genomic DNA from the mutant, as shown in **Fig. 3A**. The whole cell absorption spectrum of this mutant (**Fig. 3B**) confirmed deletion of *bciA* did not impact BChl *a* biosynthesis as the native COR*a* enzyme is able to perform C8-vinyl reduction, as expected (Canniffe et al. 2013); this mutant assembles the reaction centre–light harvesting complex 1 (RC–LH1) supercomplex and light harvesting complex 2 (LH2) antenna, absorbing maximally at 875 nm, and 800 and 850 nm, respectively. Subsequently, the *bchYZ* genes encoding the catalytic subunits of COR*a* were deleted in both the WT and Δ*bciA Rba. sphaeroides* strains to generate a background in which to determine if the *Roseilinea* COR enzyme was able to reduce the C7=C8 bond of Chlide, and to confirm that this enzyme encoded the COR*a* isoform, rather than COR*b*. Finally, the *bchNB* genes encoding the catalytic subunits of DPOR were also deleted in WT, Δ*bciA*, Δ*bchYZ* and Δ*bciA* Δ*bchYZ* strains, generating backgrounds in which to investigate whether the *Roseilinea* COR enzyme is also able to perform the C17=C18 reduction usually performed by DPOR, acting as the ancient promiscuous reductase proposed by Ward and Shih (Ward and Shih 2021). Whole cell absorption spectra demonstrated that deleting either *bchYZ* or *bchNB* abolished BChl biosynthesis, preventing assembly of LH2 and RC–LH1, and lead to an accumulation of Chlide (absorption feature at ∼670 nm) or PChlide (absorption feature at ∼630 nm) when COR and/or DPOR, respectively, were deleted, as seen in **Fig. 3B**.

**Figure 3.**
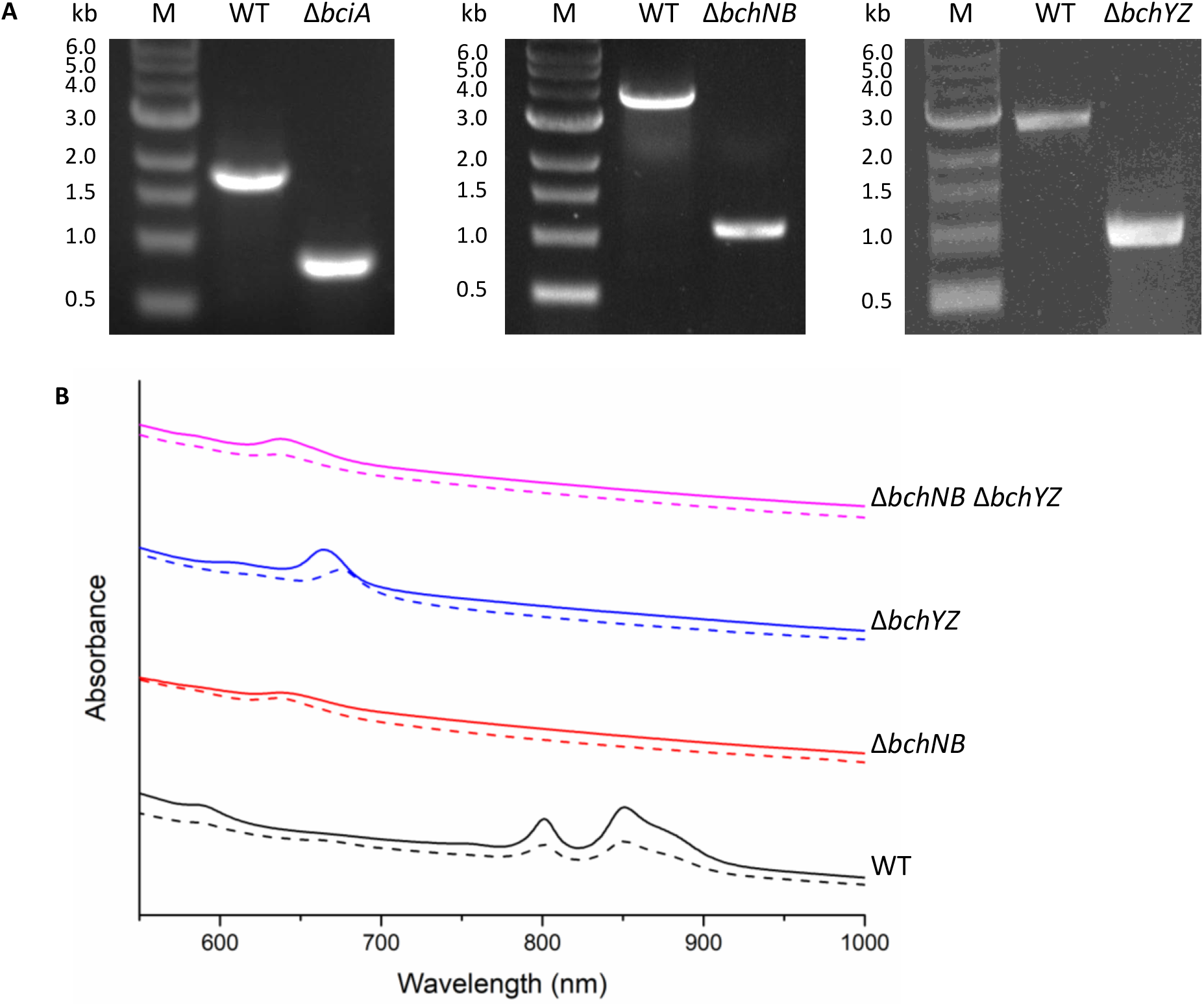
Knockout of *bciA*, *bchNB* and *bchYZ* genes from *Rba. sphaeroides*. A) PCR screening confirms gene knockouts. B) Whole cell absorption spectra show BChl biosynthesis is abolished in all mutants lacking *bchNB* or *bchYZ* as shown by the lack of B800, B850 and B875 peaks. Δ*bciA* strains are shown using dashed lines. Normalised to A_740 a_nd offset for clarity.

To ascertain whether the *Roseilinea bchYZ* genes (*bchYZ^Rl^)* encode a functional catalytic component of a COR*a* enzyme, the genes from *Roseilinea* sp. NK_OTU-006 were introduced into Δ*bchYZ* and Δ*bciA* Δ*bchYZ Rba. sphaeroides* strains on the pBBRBB-P*puf*_843-1200 e_xpression plasmid under the control of a strong, constitutive promoter. Initially, the electron-shuttling ATPase-encoding *bchX* gene from *Roseilinea* sp. NK_OTU-006 was omitted as it was assumed that the native BchX from *Rba. sphaeroides* would be sufficient to provide electrons to the catalytic subunits, as has previously been the case with recombinant COR complexes composed of subunits from different bacterial phyla (Li et al. 2023). However, the whole cell absorption spectra of the transformed strains showed *bchYZ^Rl^* expression did not restore assembly of the light-harvesting complexes found in the control strains (**Fig. 4**). A BLASTP search revealed that BchX proteins from *Rba. sphaeroides* and *Roseilinea* sp. NK_OTU-006 share only 45.6% identity, so it was thought that the FAP enzyme may function in the native organism but have reduced activity in *Rba. sphaeroides* due to the incompatibility of the native electron shuttle with the recombinant catalytic subunits. Therefore, the *Roseilinea* sp. NK_OTU-006 *bchX* gene was added to the pBBRBB[*bchYZ^Rl^*] plasmid, downstream of the *bchYZ* genes and under the control of an additional *puf* promoter. This pBBRBB[*bchYZ^Rl^-bchX^Rl^*] plasmid was introduced into the *Rba. sphaeroides* strains and whole cell absorption spectra revealed BChl *a* biosynthesis was restored in Δ*bchYZ* and Δ*bciA* Δ*bchYZ* mutants, confirming that the *bchXYZ^Rl^*genes encode a functional COR*a* enzyme (**Fig. 4**).

**Figure 4.**
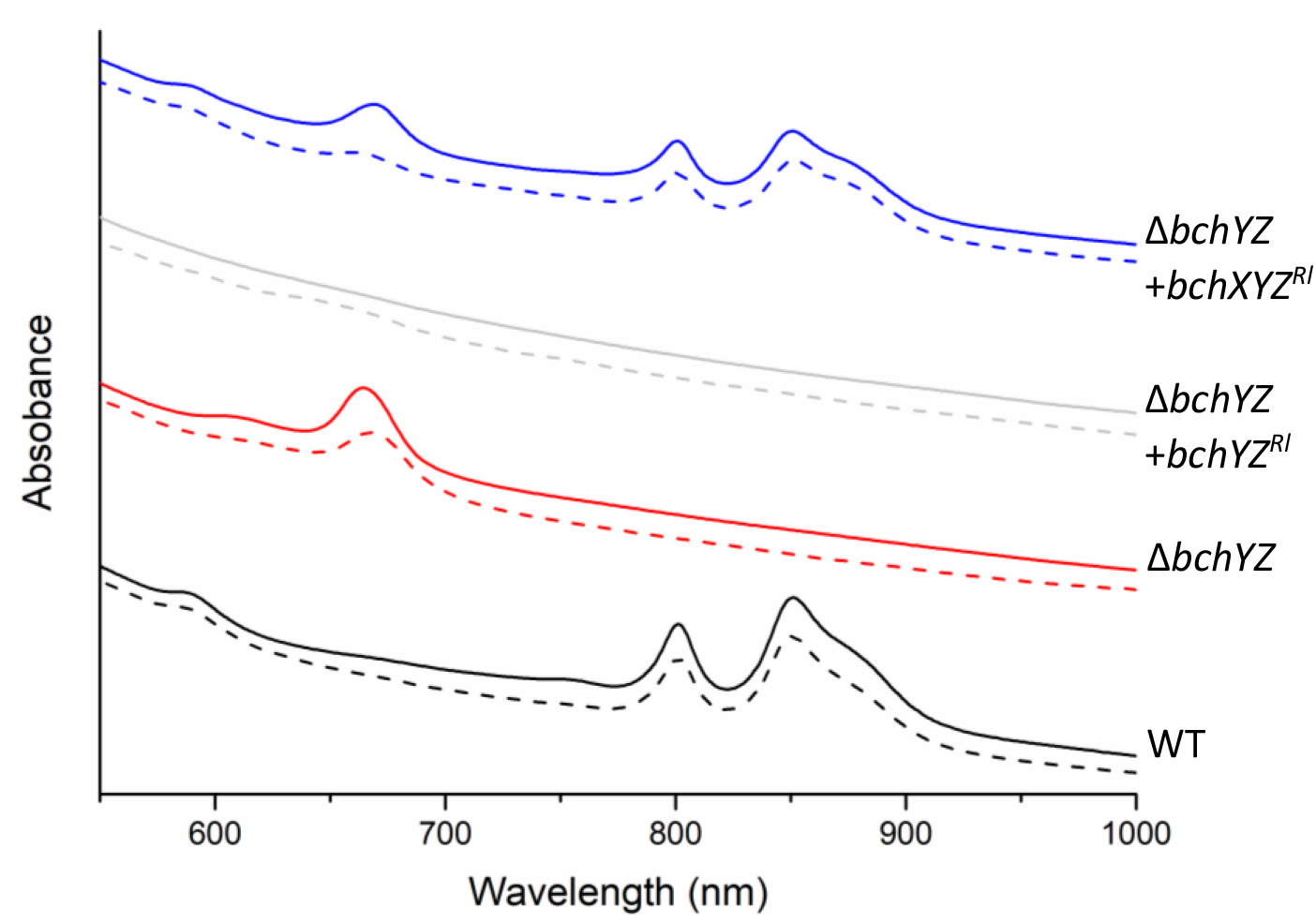
Expression of *bchYZ^Rl^*does not restore BChl biosynthesis in Δ*bchYZ Rba. sphaeroides* without *bchX^Rl^*. Whole cell absorption spectra show Δ*bchYZ* and Δ*bciA* Δ*bchYZ Rba. sphaeroides* lack the B800, B850 and B875 peaks. Background strains containing and lacking *bciA* are shown using solid and dashed lines, respectively. All traces are normalised to A_740 a_nd offset for clarity.

To determine whether the *Roseilinea* COR*a* could be an example of the hypothesised, ancestral dual-functional oxidoreductase, the pBBRBB[*bchYZ^Rl^-bchX^Rl^*] plasmid was then introduced into mutants lacking *bchNB* to investigate if the C17=C18 could also be reduced by the *Roseilinea* enzyme. However, no BChl biosynthesis was detected in any mutant lacking *bchNB* (**Fig. 5**). These mutants instead continued to accumulate PChlide, indicating the *Roseilinea* COR*a* enzyme is able to reduce the C7=C8 macrocycle bond and 8-vinyl group, but not the C17=C18 usually performed by DPOR. HPLC was performed to verify no traces of BChl were produced in the *bchNB* mutants expressing the *Roseilinea* COR genes. As shown in **Fig. 6A**, BChl *a* was seen in the WT and Δ*bciA* strains but was absent in all of the additional mutant controls. When pBBRBB[*bchYZ^Rl^-bchX^Rl^*] was conjugated into each of these strains, restoration of BChl biosynthesis was only observed in Δ*bchYZ* and Δ*bciA* Δ*bchYZ* strains but not in those lacking *bchNB* (**Fig 6B**). These results confirm that the *Roseilinea* COR enzyme has no C17=C18 reductase activity. Interestingly, this suggests *Roseilinea* spp. must either have an alternative enzyme for protochlorophyllide reduction unlike any seen in nature, which could not been identified using homology searches, or must have an alternative method of acquiring Chlide.

**Figure 5.**
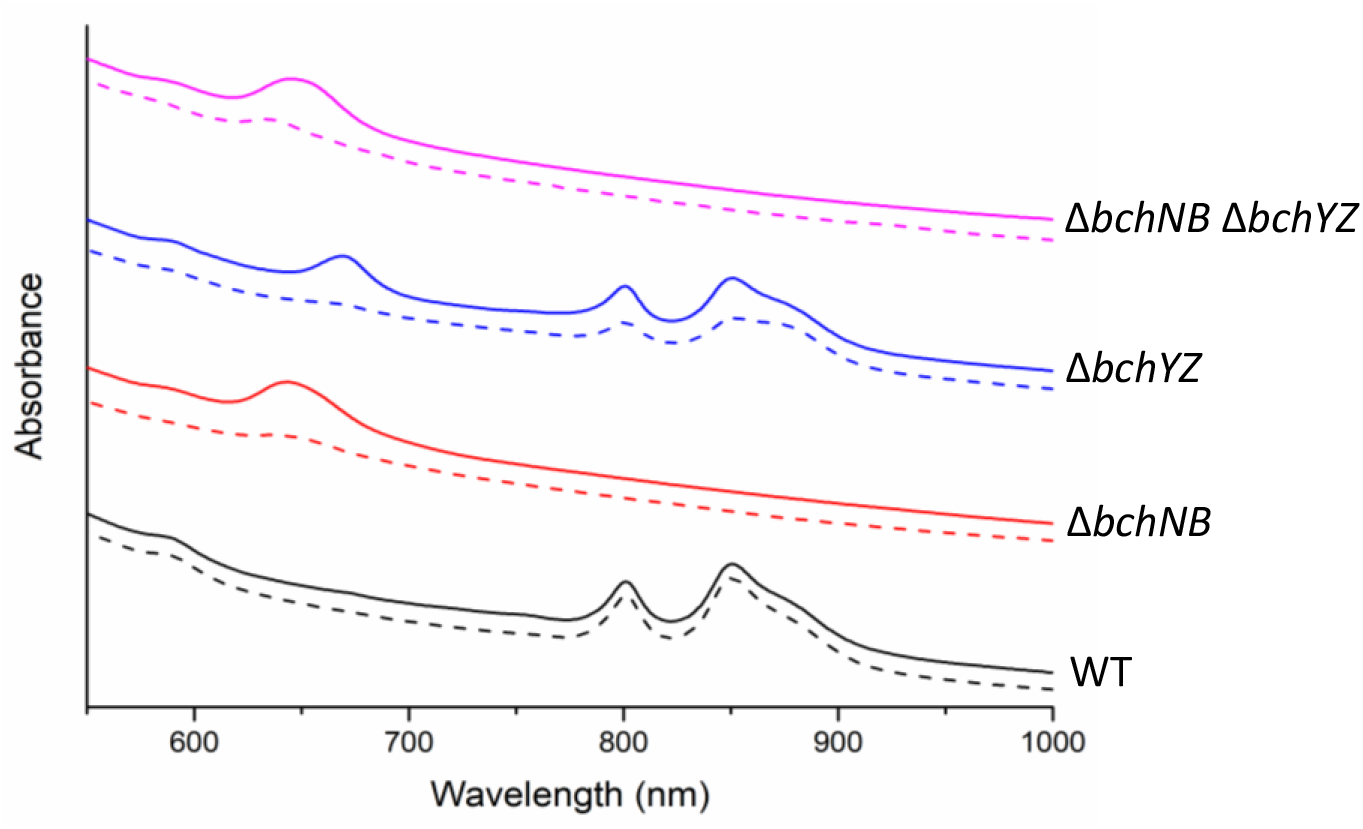
Whole cell absorption spectra of strains expressing *Roseilinea* COR. BChl biosynthesis is restored in Δ*bchYZ* and Δ*bciA* Δ*bchYZ* as shown by peaks from LH2 at 800 and 850 nm and RC–LH1 shoulder at 875 nm. Δ*bciA* strains are shown using dashed lines. Normalised to A_740 a_nd offset for clarity.

**Figure 6.**
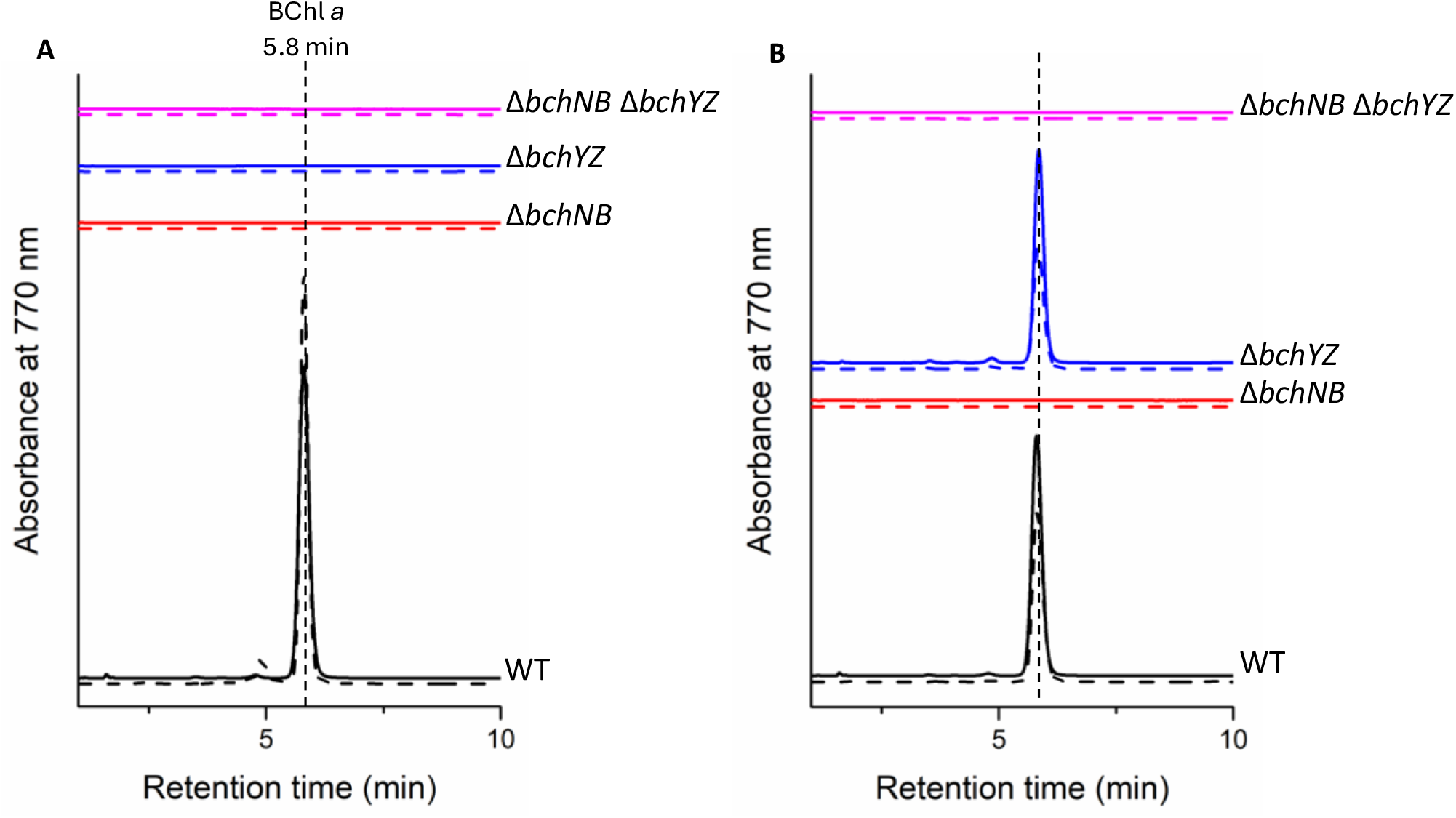
HPLC analysis of all *Rba. sphaeroides* strains in (A) the absence and (B) the presence of pBBRBB[*bchYZ^Rl^-bchX^Rl^*]. Retention times attributed to peak from BChl *a* is shown. Cultures were normalised to the same OD_740 p_rior to pigment extraction.

HPLC was also performed on supernatant from each strain in the presence and absence of pBBRBB[*bchYZ^Rl^-bchX^Rl^*] to identify any BChl precursors accumulating in each strain. **Fig. 7** shows all strains lacking *bchNB* accumulated PChlide as expected. However, despite strains lacking the native *bchYZ* being able to synthesise BChl *a* in the presence of pBBRBB[*bchYZ^Rl^-bchX^Rl^*], these cells still accumulated either 8V-Chlide or 8E-Chlide, depending on whether *bciA* was present. It is not surprising that some pigment precursors accumulated in the Δ*bchYZ* strains since the *Roseilinea* COR*a* may have reduced activity in *Rba. sphaeroides*.

**Figure 7.**
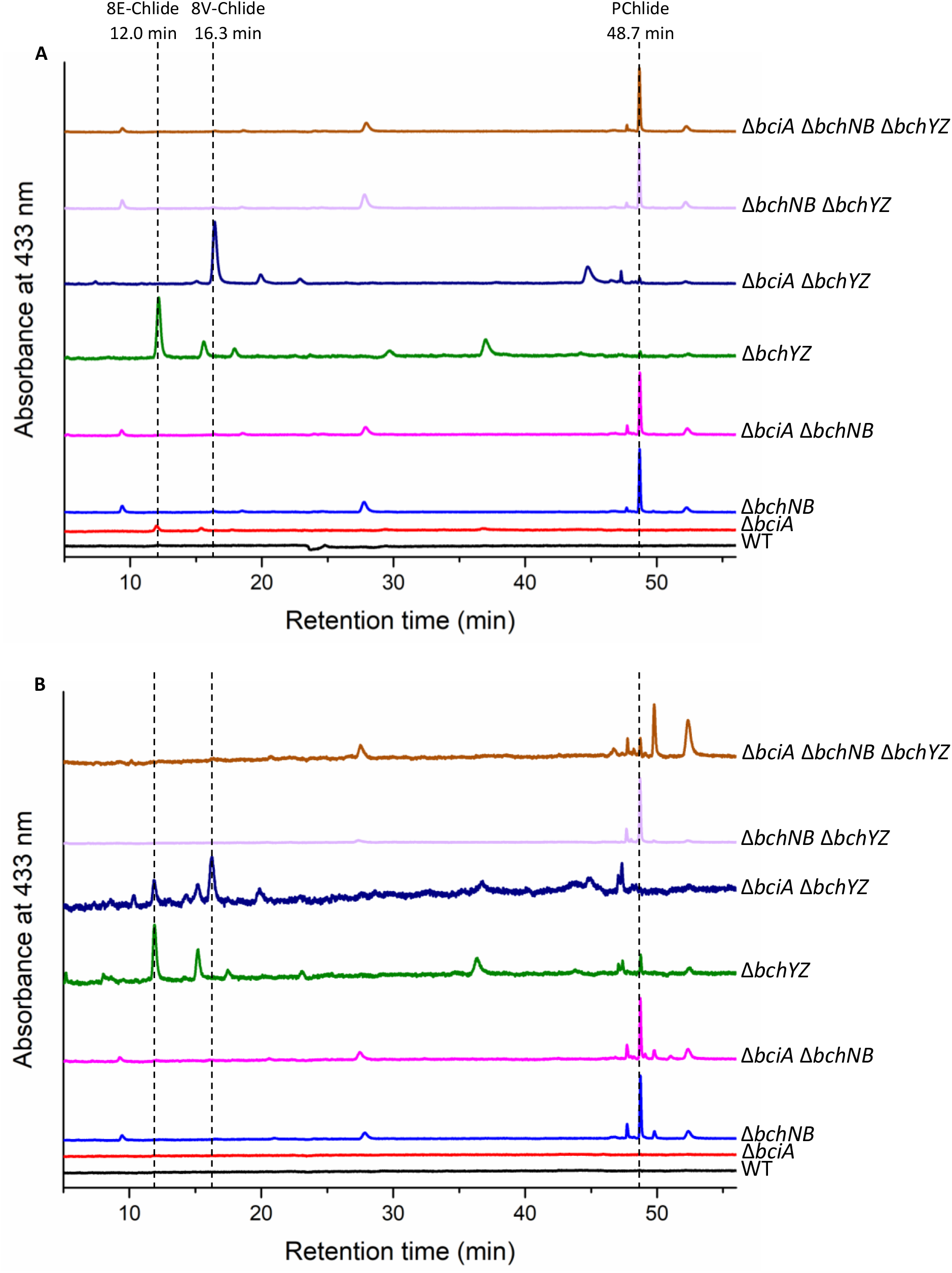
HPLC analysis of spent media from all *Rba. sphaeroides* strains in (A) the absence and (B) the presence of pBBRBB[*bchYZ^Rl^-bchX^Rl^*]. Retention times attributed to peaks from 8E-Chlide, 8V-Chlide and PChlide are shown. Normalised to maximum peak height and offset for clarity.

### Domain swapping in BchX to uncover the roles of N– and C-terminal extensions

To determine whether there was an obvious structural reason for the BchX*^Rs^* subunit being unable to provide electrons to Bch(YZ)_2*R*_*^l^*, AlphaFold models were constructed of BchX subunits from both species. These revealed an N-terminal α-helix that was present in the *Rba. sphaeroides* protein but missing in BchX*^Rl^*, and a C-terminal domain present in *Roseilinea* that was missing in the *Rba. sphaeroides* protein, shown in **Fig. 8A**. Previously, it has been shown that the *Rba. sphaeroides* electron shuttle to DPOR, BchL, contains a similar N-terminal region to that in BchX, which confers an autoinhibitory function to prevent BchL binding to Bch(NB)_2 i_n the absence of ATP (Corless et al. 2021). Although the key residues involved in this activity from BchL are not conserved in BchX, it is possible that other residues perform this activity in BchX. Therefore, to determine whether the incompatibility between BchX and the catalytic component of the two COR*a* enzymes was due to the *Rba. sphaeroides* BchX N-terminus inhibiting activity or the *Roseilinea* BchX C-terminus being essential for its interaction with Bch(YZ)_2,_ BchX*^Rs^* variants were constructed either lacking the native N-terminus or containing the *Roseilinea* C-terminus addition, and BchX*^Rl^* variants were constructed either lacking the native C-terminus or containing the *Rba. sphaeroides* N-terminus addition. All four BchX variants were introduced into Δ*bchX Rba. sphaeroides* on the pBBRBB expression plasmid to determine their ability to shuttle electrons to Bch(YZ)_2*R*_*^s^* and into Δ*bciA* Δ*bchYZ Rba. sphaeroides* on the pBBRBB expression plasmid already containing *bchYZ^Rl^* to determine their ability to provide electrons to Bch(YZ)_2*R*_*^l^*. It was found that both the ΔN BchX*^Rs^* and BchX*^Rs^* with the C*^Rl^* addition could interact with Bch(YZ)_2*R*_*^s^* and restore BChl *a* biosynthesis (**Fig. 8B**). However, of the four BchX variants constructed, only BchX*^Rl^* with the N*^Rs^* addition was able to transfer electrons to Bch(YZ)_2*R*_*^l^* to restore BChl *a* biosynthesis. This suggests that the N-terminal helix of the *Rba. sphaeroides* BchX does not have an autoinhibitory effect, in agreement with the findings of Corless and colleaugues (Corless et al. 2021).

**Figure 8.**
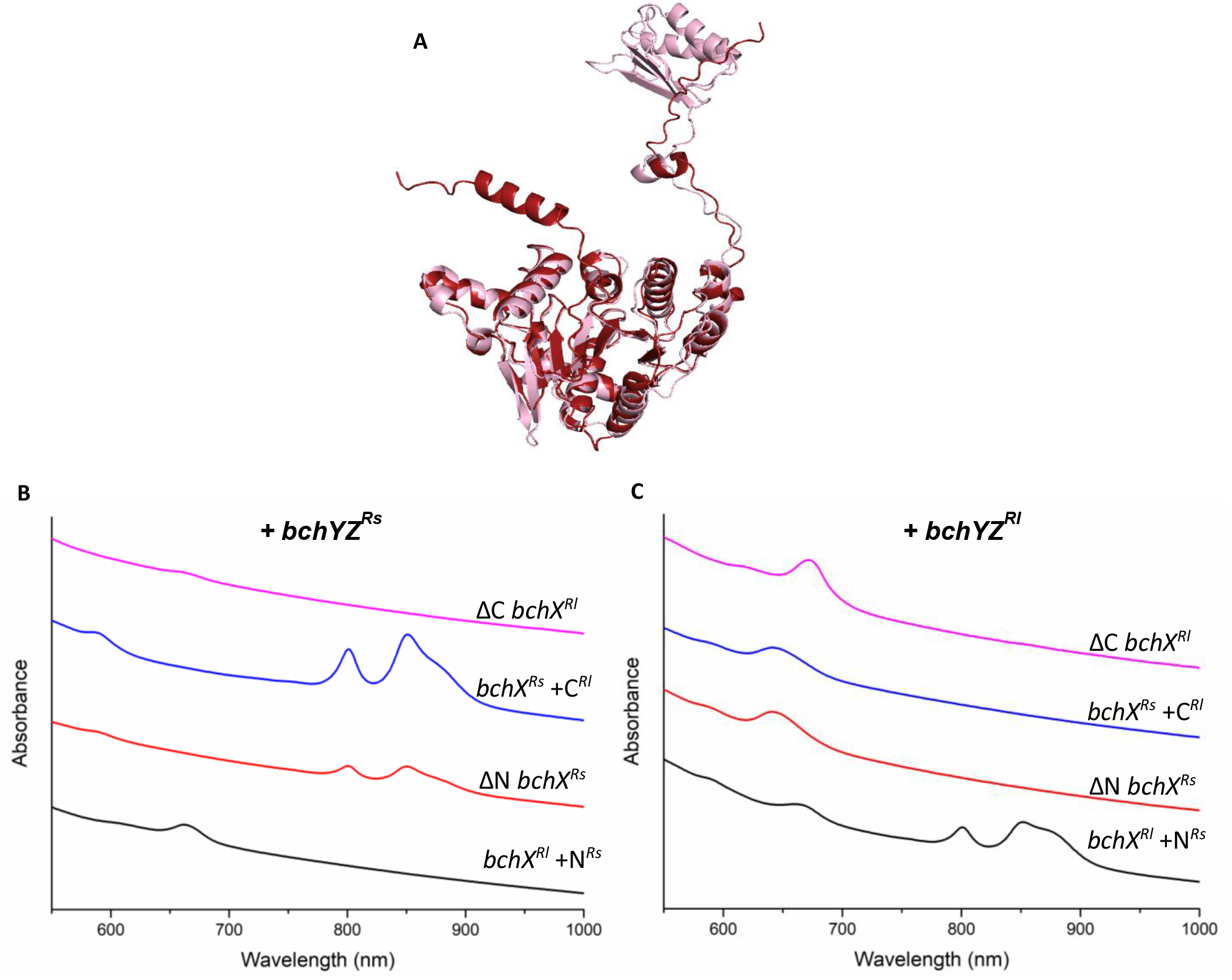
BchX subunits are not interchangeable between *Roseilinea* and *Rba. sphaeroides* COR*a* enzymes. A) AlphaFold models of BchX from *Rba. sphaeroides* (red) and *Roseilinea* sp. NK_OTU-006 (pink). B) Absorption spectra show that only BchX*^Rs^*, not BchX*^Rl^*, is able to transfer electrons to BchYZ*^Rs^*, however the N-terminal helix of BchX*^Rs^* is not required for electron transfer. C) Absorption spectra show that only BchX*^Rl^*, not BchX*^Rs^*, is able to transfer electrons to BchYZ*^Rl^*, but addition of the N-terminal helix of BchX*^Rs^* did not inhibit electron transfer.

### Reconstruction of the terminal steps of *Roseilinea* BChl biosynthesis

Since *Roseilinea* lack the central enzymes of the BChl biosynthesis pathway, while the documented fluorescence emission of these cells indicates their ability to synthesise these pigments (Tank et al. 2017), we sought to confirm that the entire late-stage BChl biosynthesis pathway encoded in “*Ca*. Roseilinea sp. NK_OTU-006” is functional. To do this, the *bchF*, *bchC*, *bchG* and *bchP* genes from this strain were synthesised and again cloned into the pBBRBB plasmid under the control of the *puf* promoter and introduced into *Rba. sphaeroides* mutants lacking the equivalent native gene. Whole cell absorption spectra revealed that the *Roseilinea bchF* and *bchC* genes restored BChl biosynthesis to some extent in the respective *Rba. Sphaeroides* mutants, as shown in **Fig. 9A**. However, the *Roseilinea bchG* did not appear to restore BChl biosynthesis, as seen by the lack of the B800 and B850 peaks in the whole cell absorption spectrum. The *Roseilinea* genes are not closely related to those in purple bacteria and their encoded proteins may have lower activity in *Rba. sphaeroides*, which would explain the reduced BChl biosynthesis. Therefore, the mutants were incubated under phototrophic conditions to determine whether the *Roseilinea* genes were able to support phototrophic growth. The strains expressing the *Roseilinea bchF* and *bchC* genes were able to grow under these conditions, but the heterologous expression of *bchG* did not support phototrophic growth, as seen in **Fig. 9B**. Complementation by the *Roseilinea bchP* gene could not be investigated by whole cell absorption spectroscopy as reduction of the BChl geranylgeranyl tail by BchP does not affect the absorption properties of the pigment. Therefore, HPLC was performed to study the function of BchP^Rl^. As shown in **Fig. 9B**, in the Δ*bchP* mutant, the retention time of BChl *a*_GG w_as shorter than in the WT containing BChl *a* with a phytyl tail; in the presence of the *Roseilinea bchP* the WT peak is restored, demonstrating that BchP from *Roseilinea* is able to reduce the BChl *a* geranylgeranyl tail to a phytyl tail.

**Figure 9.**
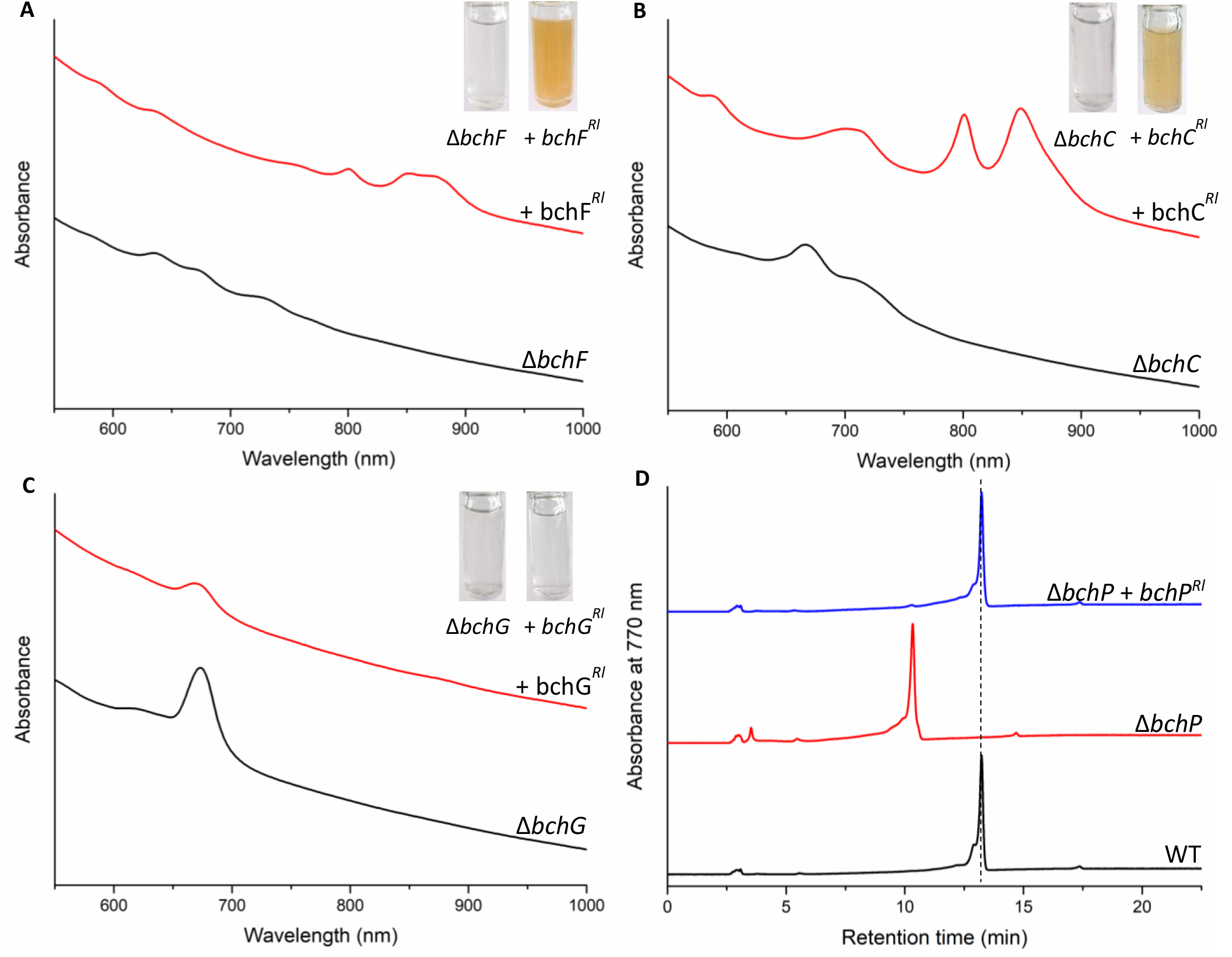
*bchF^Rl^* and *bchC^Rl^* and *bchP^Rl^* expression allow BChl biosynthesis and phototrophic growth in *Rba. sphaeroides*. A) Absorption spectra of Δ*bchF*, B) Δ*bchC* and C) Δ*bchG* in the absence (black) and presence (red) of a *Roseilinea* complementation plasmid. BChl biosynthesis is restored in Δ*bchF* and Δ*bchC* as shown by peaks from LH2 at 800 and 850 nm and RC–LH1 shoulder at 875 nm. Spectra are normalised to A_740 a_nd offset for clarity. Inset photos show phototrophically-grown cultures of *Rba. sphaeroides* mutants with and without complementation from *Roseilinea* genes. *bchF^Rl^* and *bchC^Rl^* support phototrophic growth whereas *bchG^Rl^* does not. D) HPLC analysis of Δ*bchP* in the absence and presence of the *Roseilinea* complementation plasmid shows BChl tail reduction is restored by *bchP^Rl^*.

It was thought that the lack of BchG activity may be due to overproduction of this integral membrane protein being detrimental to cell growth when using a strong promoter and a multi-copy replicating plasmid. Therefore, the *Roseilinea bchG* was integrated into the genome of *Rba. sphaeroides* in place of the native *bchG*, as shown in **Fig. 10A**, allowing a lower level of expression under control of the native promoter. Whole cell absorption spectra revealed the integrated *Roseilinea bchG* produced a small RC–LH1 875 nm peak, indicating this enzyme is active in BChl *a* biosynthesis (**Fig. 10B**). Additionally, it was found that this mutant was capable of growing phototrophically, unlike when *bchG^Rl^ was* introduced on an expression plasmid, as seen in **Fig. 10C**. Under phototrophic conditions the Δ*bchG::bchG^Rl^* mutant showed peaks at 800 and 850 nm demonstrating that BChl *a* could be synthesised at a level to permit assembly of LH2, as well as RC–LH1. The integrated *bchG^Rl^* gene was sequenced to confirm the activity was not due to the introduction of a gain-of-function mutation. No mutations were seen in *bchG^Rl^*, confirming the encoded *Roseilinea* enzyme is functional when expressed from the genome of *Rba. sphaeroides* under the control of the native promoter.

**Figure 10.**
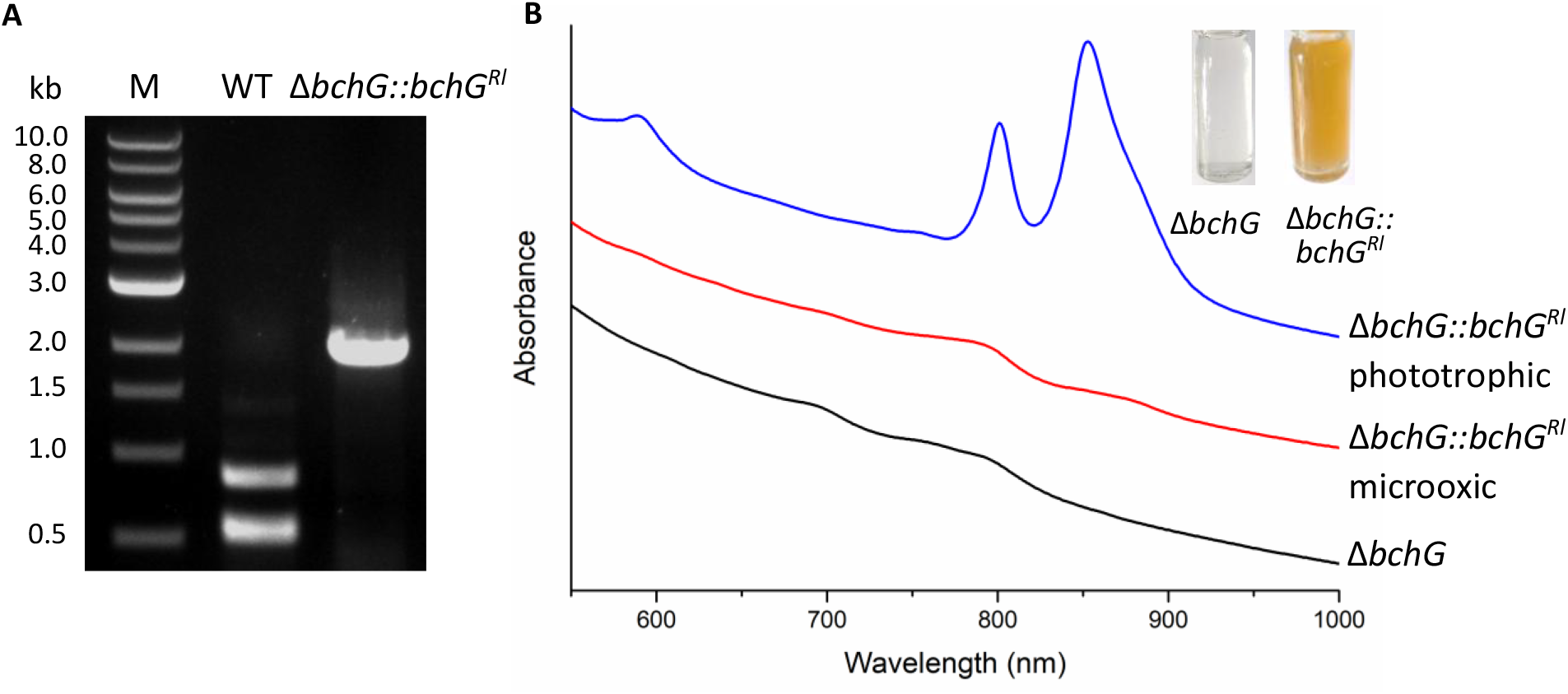
Genomic integration of *bchG^Rl^* restores BChl biosynthesis in Δ*bchG Rba. sphaeroides*. (A) PCR screening confirms replacement of *bchG* from *Rba. sphaeroides* with that from *Roseilinea* sp. NK_OTU-006. (B) Whole cell absorption spectrum shows B875 peak is restored in the *bchG^Rl^* mutant. Inset photos show the *bchG^Rl^*mutant is capable of phototrophic growth.

### Co-culturing of BChl biosynthesis mutants reveals a precursor import mechanism

To determine whether it is possible for soluble BChl precursors to be exchanged between bacteria in a microbial community, the Δ*bchNB* mutant was co-cultured with a Δ*bchE* mutant of *Rba. sphaeroides* (Chen et al. 2016) in liquid media. BchE is required to convert Mg-protoporphyrin IX monomethyl ester (MgPME) to 8V-PChlide; the Δ*bchE* mutant accumulates MgPME and is unable to synthesise BChl *a* (Chen et al. 2016). Since PChlide is excreted from Δ*bchNB Rba. sphaeroides*, which is also unable to make BChl *a*, this pigment may be taken up by Δ*bchE* cells, and the native terminal enzymes able to act on this exogenous metabolite to restore BChl biosynthesis. As shown in **Fig. 11**, a small absorption feature is seen at approximately 875 nm when the two strains are co-cultured, which is not present when the strains are grown individually. This implies PChlide can be imported by Δ*bchE*, permitting BChl *a* synthesis and assembly of RC–LH1. This precursor import to restore BChl biosynthesis is confirmed by HPLC analysis of these three strains, with only the co-culture extract containing BChl *a* (**Fig 11.B**)

**Figure 11.**
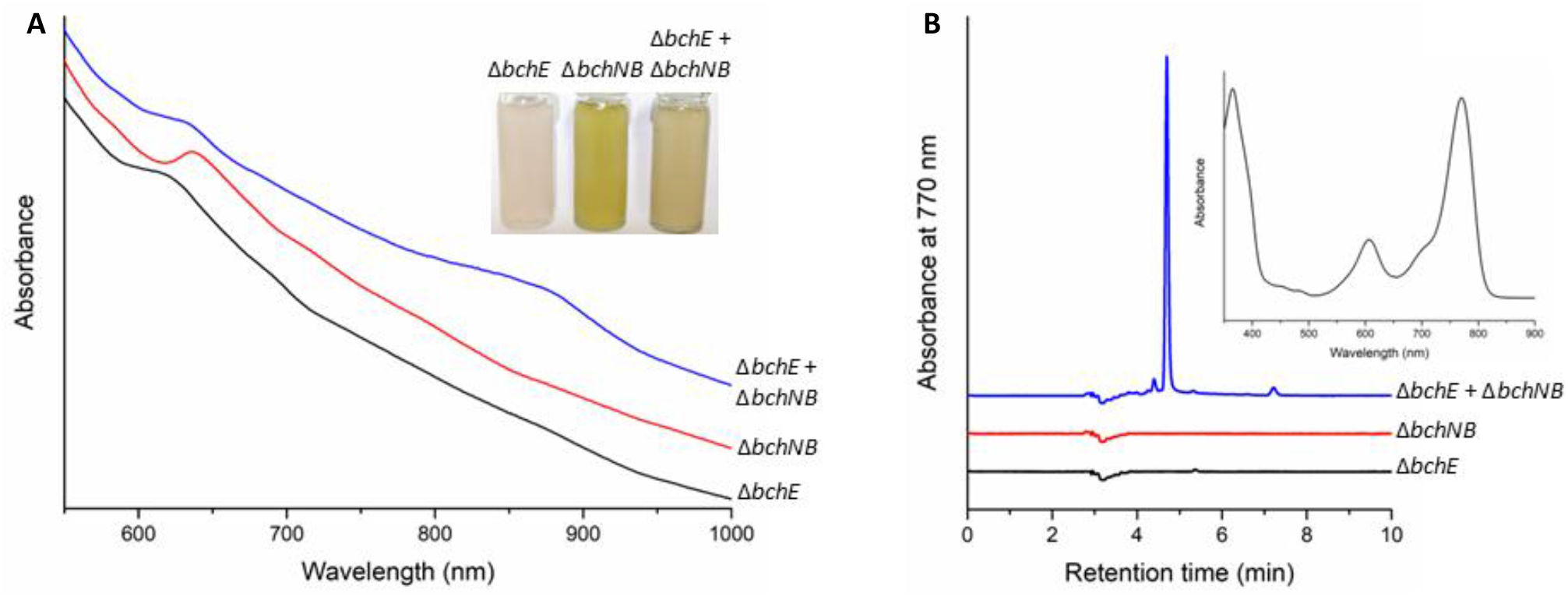
Co-culture of Δ*bchE Rba. sphaeroides* and Δ*bchNB Rba. sphaeroides* shows BChl precursors can be shared between strains. (A) Whole cell absorption spectra of bch mutants, grown under microoxic conditions alone, and in co-culture. Inset shows pigmentation of the individual and co-cultured mutants. (B) HPLC analysis of culture extracts. Inset shows the spectrum of the peak at 4.7 min, corresponding to BChl *a*.

## DISCUSSION

Our results confirm *Roseilinea* spp. contain a functional COR*a* enzyme capable of reducing the C7=C8 double bond and 8-vinyl group on Chlide, a prerequisite for the biosynthesis of BChl *a*. In addition, we have shown that the terminal enzymes BchF, BchC, BchG, and BchP from *Ca*. Roseilinea sp. NK_OTU-006 are functional and able to restore BChl biosynthesis in *Rba. sphaeroides* mutants lacking the native genes. Although overexpression of *bchG^Rl^* did not restore BChl biosynthesis, genomic integration of this gene was able to support phototrophic growth. Since BchG is an integral membrane protein with multiple transmembrane helices (Proctor et al. 2022), overproduction may have a negative impact on the cells by crowding the membrane, or its insertion into a non-native membrane—perhaps containing different lipids and/or having a different bilayer thickness (Pogozheva et al. 2013)—may have contributed to the failure to detect activity from plasmid-bourne expression (Jensen et al. 2017).

Most BChl *a*-dependent phototrophs contain genes encoding canonical BciA and/or BciB classes of C8-vinyl reductase, despite having the COR*a* isoform able to perform this same reaction. However, *Roseiflexus* spp., chlorosome-lacking FAPs like *Roseilinea*, are one such group that relies solely on COR*a* for C8-vinyl reduction in the synthesis of BChl *a*. The absence of *bciA*/*bciB* in *Roseiflexus* and *Roseilinea* spp. may imply that the common ancestor of the two classes lost one of these genes. Alternatively, the common ancestor may have diverged from the chlorosome-containing *Chloroflexia* before the latter acquired *bciA*/*bciB*, and may be a key determining factor for *Chloroflexia* utilizing chlorosomes; deletion of *bciA* in the green sulfur bacterium *Chlorobaculum tepidum* resulted in the formation of chlorosome pigments unable to be methylated at the C8 position by BchQ, decreased synthesis of total chlorosome pigments, blue-shifted chlorosome absorption, and decreased the half-bandwidth of Q_y a_bsorption (Thweatt et al. 2019).

Since phototrophic *Roseilinea* lack not only a POR enzyme (BchLNB or LPOR) but also BchM and BchE, they may have an alternative, unidentified pathway for synthesising BChl using novel enzymes. Previously, the probability of *bchLNB*, *bchM* and *bchE* being present in *Roseilinea* but not identified from metagenomic data was calculated to be <0.005% (Ward et al. 2018), therefore it is extremely unlikely that a conventional DPOR enzyme is present. *Roseilinea* also lack the unrelated light-activated POR enzyme utilised by oxygenic phototrophs, so must either contain a POR enzyme unlike any others seen in nature or must have an alternative method of producing BChl.

As phototrophic, autofluorescent *Roseilinea* are found in microbial mats with many other phototrophic species, the possibility exists that they are able to scavenge pigments from their surroundings to use for phototrophic growth. It was recently discovered that a non-phototrophic member of the closely related *Anaerolineae*, recovered in a MAG from an estuary sediment (Baker et al. 2015), encodes an enzyme able to remove the central Mg ion from Chl *a*, the first step in the Chl breakdown pathway visible in deciduous trees in autumn (Dey et al. 2021). This enzyme had expanded activity when compared to its plant orthologue, dechelating Mg from Chl *b* and the algal antenna Chl *c* species (Obata et al. 2019). It is possible that this organism uses this enzyme in a pathway to scavenge carbon and nitrogen from Chl macrocycles and as an acquisition mechanism for Mg^2+^. The synthesis of (B)Chls is metabolically and energetically costly, e.g. the insertion of the central metal by Mg chelatase requires the hydrolysis of 15–40 molecules of ATP (Adams et al. 2020); the ability to scavenge late-stage (B)Chl intermediates from the environment would allow the cell to divert carbon and nitrogen to other metabolites and liberate ATP and reductant for other cellular processes. Many bacteria are auxotrophs for vitamin B12, a complex modified tetrapyrrole that acts as a cofactor in methyltransferase enzymes (e.g. the MetH methionine synthase); these organisms must acquire vitamin B12 from their environment, and actively transport this molecule into the cell. Similarly, many *Bacteroidota* are heme auxotrophs. One such organism, *Porphyromonas gingivalis*, lacks many heme biosynthesis genes but encodes four late-stage enzymes, including ferrochelatase, the final enzyme of the canonical pathway that inserts iron into protoporphyrin IX (Śmiga et al. 2026). This bacterium can grow in medium supplemented with either heme or protoporphyrin IX, suggesting that the terminal enzymes are expressed and active, and that loss of the central pathway is an evolutionary adaptation to the availability of heme and its precursors in its environment. It is possible that the phototrophic *Roseilinea* take a similar approach to acquire modified tetrapyrroles as cofactors for photosynthesis. These organisms are members of microbial mats, where cell densities are incredibly high, e.g. the concentration of cyanobacteria in the top 2 mm of a microbial mat is ten times that found in the top 200 m of ocean (Ferris et al. 1996). Death and lysis of phototrophic members of the mats will be continuous (Zorz et al. 2023), likely releasing exogenous tetrapyrrole molecules; we hypothesise that the phototrophic *Roseilinea* use this release of biomass to provide precursor molecules as substrates for the terminal BChl biosynthesis enzymes. This scenario may partly explain why these organisms have not been isolated into axenic culture in the laboratory; we propose that future isolation experiments should be performed with the supplementation of (B)Chlide in the growth medium. To this end, *Roseilinea* sp. NK_OTU-006 encodes putative transporters for heme (WP_240550393) and vitamin B12 (WP_240550448.1 and WP_320447966.1); it is possible that one of these systems could also import structurally similar (B)Chls/(B)Chlides, or they could employ an as yet unidentified transporter that is specific to (bacterio)chlorin molecules. Despite our co-culture experiment being performed on a model species outside the *Chloroflexota* phylum, the result indicates that (B)Chl intermediates can be exchanged between cells to complete the biosynthesis of mature pigments using exogenously acquired precursors.

If *Roseilinea* spp. must scavenge (B)Chls/(B)Chlides from their environment to be able to assemble their light-harvesting apparatus, an alternative mode of growth until phototrophy has been established would be beneficial to these cells. The genome of Roseilinea sp. NK_OTU-006 contains operons for both *aa*_3-_ and *cbb*_3-_type cytochrome c oxidases, indicating that it can grow by aerobic respiration, a metabolic mode that *Chloroflexus* and *Roseiflexus* spp. are capable of using (Madigan et al. 1974; Klatt et al. 2013b).

Despite the work we present here being insufficient to demonstrate the existence of the proposed dual-functional reductase enzyme able to generate BChlide from PChlide, our work does not rule out the BChl-first hypothesis, which argues that Chl-containing photosystems are more complex than the BChl dependent RCs of anoxygenic phototrophs. BChl *g*, found in the primitive homodimeric heliobacterial RC, is structurally similar to Chl *a.* It is possible that an ancestor of this photosystem operated using a Chl pigment, and the reduction of the C7=C8 group of this pigment, catalysed by a newly-evolved COR enzyme, allowed the organism to access an unused spectral window, with minimal modification of the RC protein scaffold required to house the new BChl and remain functional. Conversely, BChl *g* (and BChl *b*) carries an ethylidene group at C8 that can readily isomerise to generate the chlorin C7=C8; it has been suggested that this was the evolutionary route to Chl biosynthesis (Kobayashi et al. 1998). Supporting this, an analysis of DPOR, COR and nitrogenase subunits indicated that COR subunits are ancestral to DPOR, and that BChl-based anoxygenic photosynthesis originated prior to the chlorophyll (Gupta 2012). Although it is unlikely that the proposed dual-functional reductase will be identified in an extant phototroph, the possibility of designing a promiscuous ‘ancestral’ enzyme now exists (Prakinee et al. 2024), which—if successful—will add further weigh to the BChl-first hypothesis.

## MATERIALS & METHODS

### Growth of described strains

*Rba. sphaeroides* strains (constructed in the 2.4.1 parental strain) were grown under microoxic conditions in the dark at 30°C in liquid M22+ medium (Hunter and Turner 1988) formulated with 50% of the original nitrilotriacetic acid content, and supplemented with 0.1% (w/v) casamino acids in a rotary shaker, or under phototrophic conditions in glass tubes filled with anoxic sodium succinate medium (Claus and Schaub-Engels 1977) on a rotating wheel, under illumination with 100W incandescent bulbs.

Transformed *E*. *coli* strains NEB5α (New England Biolabs) and ST18 (Thoma and Schobert 2009) were grown in a rotary shaker at 37°C in LB medium supplemented with 30 μg·ml^-1^ kanamycin. ST18 cells were supplemented with 50 μM δ-aminolaevulinic acid. All strains and plasmids used in the present study are listed in **Supplementary Table S1**.

### Co-culture of *Rba. sphaeroides* mutants

Starter cultures of each *Rba. sphaeroides* strain were grown in liquid modified M22+ medium overnight under microoxic conditions in the dark. 1 ml of each culture was used to co-inoculate a fresh culture and grown overnight as before. Cell pellets were collected by centrifugation (6500 x *g* for 10 min) and resuspended in 50% sucrose solution (w/v) for whole cell absorption analysis.

### Construction of deletion mutants of *Rba. sphaeroides*

*Rba. sphaeroides* genes were deleted using the allelic exchange vector pK18*mobsacB* **(Schäfer et al. 1994)**. Sequences up– and down-stream of target genes were amplified with the relevant UpF and UpR, and DownF and DownR primers respectively. Sequences of the primers used in the present study can be found in **Supplementary Table S2**. The up– and down-stream PCR products were fused by overlap-extension PCR, and the final amplicon was digested with relevant restriction enzymes and ligated into cut pK18*mobsacB*. Sequenced clones were conjugated into *Rba. sphaeroides* from *E. coli* ST18, and transconjugants in which the clone had integrated into the genome by homologous recombination were selected on modified M22+ medium supplemented with kanamycin, and lacking δ-aminolaevulinic acid to block growth of the ST18 cells. Transconjugants that had undergone a second recombination event were then selected on M22+ supplemented with 10% (w/v) sucrose, lacking kanamycin. Sucrose-resistant and kanamycin-sensitive colonies had excised the allelic exchange vector through the second recombination event. Deletion of desired genes was confirmed by colony PCR using relevant CheckF and CheckR primers.

### Expression of Roseilinea genes in Rba. sphaeroides

Gene sequences from “*Ca*. Roseilinea sp. strain NK_OTU-006” were recovered through Blast searches using the following genes from the model FAP *Chloroflexus aurantiacus* J-10-fl^T^ as search queries: *bchX* (Caur_0417), *bchY* (Caur_3805), *bchZ* (Caur_3806), *bchF* (Caur_0415), *bchC* (Caur_0416), *bchG* (Caur_2088), *bchP* (Caur_2087). The recovered sequences were optimised for expression in *Rba. sphaeroides* using a Codon Optimization tool (idtdna.com) and ordered as gBlocks from Integrated DNA Technologies, flanked with BglII and SpeI restriction sites. A fragment containing the adjacent *bchY* (WP_169238029) and *bchZ* (WP_169238028) was ordered as a single gBlock; the intergenic region between the two optimised open reading frames was also manually adjusted to contain a *Rba. sphaeroides* ribosome binding site for *bchZ*. All sequences are detailed in **Supplementary Table S3**. Restriction digested fragments were cloned into the BglII/SpeI sites of pBBRBB-*Ppuf*_843–1200 (_Tikh et al. 2014). Plasmid constructs were sequence verified with Plasmidsaurus Whole Plasmid Sequencing (Oxford Nanopore, R10.4.1). Sequenced plasmids were conjugated into relevant strains from *E. coli* ST18 and transconjugants were selected on modified M22+ medium supplemented with kanamycin. Plasmid pBBRBB-*Ppuf_843_*_–*1200* c_ontains a truncated version of the native *Rba. sphaeroides puf* promoter providing constitutive expression under oxic and anoxic growth regimes.

### Absorption spectroscopy

UV/vis/near-IR absorption spectra were collected on a Cary 3500 spectrophotometer (Agilent Technologies) scanning between 300 and 1100 nm at 1 nm intervals with a 0.1 s integration time.

### Extraction of pigments

Pigments were extracted from cell pellets after washing in 20 mM HEPES pH 7.2 by adding 9 pellet volumes of 0.2% (v/v) ammonia in methanol, vortex-mixing for 30 s and incubating on ice for 20 min. The extracts were clarified by centrifugation (15000 x *g* for 5 min at 4°C) and the supernatants were immediately analysed on an Agilent 1260 HPLC system. Carotenoids were removed from clarified extracts by the addition of 50 µl of 5 M NaCl and 9 pellet volumes of hexane. The samples were mixed and allowed to partition, the upper hexane phase was discarded and the lower methanol phase was clarified before analysis.

### Analysis of pigments by HPLC

PChlide and Chlide species were separated on a YMC30 C30 reverse-phase column (3 μm particle size, 250 mm×4.6 mm) as previously described (Canniffe et al. 2014). Solvents A and B were 10:90 (v/v) methanol/500 mM ammonium acetate, and methanol respectively. Pigments were eluted at 1 ml/min at 40°C on a linear gradient of 82–98% solvent B over 40 min, increasing to 100% to wash the column. Elution of PChlide and Chlide species was monitored by checking absorbance at 632 nm and 665 nm, respectively.

BChl species were separated on a Supelco Discovery C18 reverse-phase column (5 μm particle size, 250 mm × 4.6 mm) using a previously described method (Ortega-Ramos et al. 2018). Solvents A and B were 64:16:20 (v/v/v) methanol:acetone:H_2O_ and 80:20 (v/v) methanol:acetone, respectively. After 2 min held at 50% solvent B, pigments were eluted over a linear gradient to 100% solvent B over 10 min, held at 100% solvent B for 25 min. Elution of BChl species was monitored by checking the absorbance at 770 nm.

## AUTHOR CONTRIBUTIONS

DPC and VT conceived the study. NMR and DN conducted experiments. All authors interpreted data. DPC and NMR wrote the manuscript with contributions from DN and VT.

## RESEARCH FUNDING

NMR and DPC are supported by Biotechnology and Biological Sciences Research Council (BBSRC) grant BB/X015955/1. NMR acknowledges support from a Doctoral Training Partnership from BBSRC (BB/T008695/1). DPC acknowledges additional support from BBSRC grants BB/W008076/1 and UKRI2820. D.N. was supported by a Mahidol–Liverpool PhD Scholarship.

## DATA AVILABILITY

The authors declare that the data supporting the findings of this study are available within the paper and its Supplementary Information files. Raw data files and materials are available from the corresponding author upon reasonable request.

## ETHICS DECLARATION

Not applicable

