## Supplementary Material for "Unravelling bacteriochlorophyll biosynthesis in the uncultured *Roseilinea*"

**Supplementary tables & figures**

| Strain/Plasmid | Properties | Source/Reference |
| --- | --- | --- |
| *E. coli* |  |  |
| NEB^®^ 5-alpha | Cloning strain for pRL277 constructs | New England Biolabs |
| ST18 | Donor strain for biparental mating, Δ*hemA* in S17-1λ*pir* | DSMZ [1] |
| *Rba. sphaeroides* |  |  |
| WT | 2.4.1 | S. Kaplan† |
| Δ*bciA* | Markerless deletion of rsp_3070 in WT | [2] |
| Δ*bchYZ* | Markerless deletion of rsp_0260+rsp_0261 in WT | [3] |
| Δ*bciA* Δ*bchYZ* | Markerless deletion of rsp_0260+rsp_0261 in Δ*bciA* | [3] |
| Δ*bchNB* | Markerless deletion of rsp_0285+rsp_0286 in WT | This study |
| Δ*bciA* Δ*bchNB* | Markerless deletion of rsp_0285+rsp_0286 in Δ*bciA* | This study |
| Δ*bchNB* Δ*bchYZ* | Markerless deletion of rsp_0285+rsp_0286 in Δ*bchYZ* | This study |
| Δ*bciA* Δ*bchNB* Δ*bchYZ* | Markerless deletion of rsp_0285+rsp_0286 in Δ*bciA* Δ*bchYZ* | This study |
| Δ*bchX* | Markerless deletion of rsp_0262 in WT | This study |
| Δ*bchF* | Markerless deletion of rsp_0284 in WT | [4] |
| Δ*bchC* | Markerless deletion of rsp_0263 in WT | This study |
| Δ*bchG* | Markerless deletion of rsp_0279 in WT | [5] |
| Δ*bchG*::*bchG^Rl^* | Genomic replacement of rsp_0279 with its orthologue from *Roseilinea* sp. NK_OTU-006 | This study |
| Δ*bchP* | Markerless deletion of rsp_0277 in WT | [5] |
| Δ*bchE* | Markerless deletion of rsp_0281 in WT | [6] |
| Plasmid |  |  |
| pK18*mobsacB* | Allelic exchange suicide vector; *Km^R^* | J. Armitage‡ [7] |
| pK18*mobsacB*[*bchYZ* KO] | ~600 bp Up- and downstream of rsp_0260+rsp_0261 cloned into pK18*mobsacB* | [3] |
| pK18*mobsacB*[*bchX* KO] | ~600 bp Up- and downstream of rsp_0262 cloned into pK18*mobsacB* | This study |
| pK18*mobsacB*[*bchNB* KO] | ~600 bp Up- and downstream of rsp_0285+rsp_0286 cloned into pK18*mobsacB* | This study |
| pK18*mobsacB*[*bchC* KO] | ~600 bp Up- and downstream of rsp_0263 cloned into pK18*mobsacB* | This study |
| pK18*mobsacB*[*bchG*::*bchG^Rl^*] | Codon-optimised *bchG^Rl^* cloned between flaking regions of rsp_0279 in pK18*mobsacB* | This study |
| pBBRBB-Ppuf_843-1200_ | Expression vector containing the strong *puf* promoter of *Rba. sphaeroides*; *Km^R^* | Addgene [8] |
| pBBRBB[*bchYZ^Rl^*] | Codon-optimised *Roseilinea* sp. NK_OTU-006 *bchYZ* cloned into pBBRBB-Ppuf_843-1200_ | This study |
| pBBRBB[*bchX^Rl^*] | Codon-optimised *Roseilinea* sp. NK_OTU-006 *bchX* cloned into pBBRBB-Ppuf_843-1200_ | This study |
| pBBRBB[*bchYZ^Rl^*–*bchX^Rl^*] | Ppuf_843-1200_[*bchX^Rl^*] cloned behind *bchYZ^Rl^* in pBBRBB[*bchYZ^Rl^*] | This study |
| pBBRBB[*bchF^Rl^*] | Codon-optimised *Roseilinea* sp. NK_OTU-006 *bchF* cloned into pBBRBB-Ppuf_843-1200_ | This study |
| pBBRBB[*bchC^Rl^*] | Codon-optimised *Roseilinea* sp. NK_OTU-006 *bchC* cloned into pBBRBB-Ppuf_843-1200_ | This study |
| pBBRBB[*bchG^Rl^*] | Codon-optimised *Roseilinea* sp. NK_OTU-006 *bchG* cloned into pBBRBB-Ppuf_843-1200_ | This study |
| pBBRBB[*bchP^Rl^*]  pBBRBB[*bchYZ^Rl^*–*bchX^Rl^*+N*^Rs^*]  pBBRBB[*bchYZ^Rl^*–ΔN *bchX^Rs^*]  pBBRBB[*bchYZ^Rl^*–*bchX^Rs^*+C*^Rl^*]  pBBRBB[*bchYZ^Rl^*– ΔC *bchX^Rl^*] | Codon-optimised *Roseilinea* sp. NK_OTU-006 *bchP* cloned into pBBRBB-Ppuf_843-1200_  Ppuf_843-1200_[*bchX^Rl^*] with first 108 bp of *bchX^Rs^* fused at N-terminus cloned behind *bchYZ^Rl^* in pBBRBB[*bchYZ^Rl^*]  Ppuf_843-1200_[*bchX^Rs^*] lacking 108 bp N-terminus cloned behind *bchYZ^Rl^* in pBBRBB[*bchYZ^Rl^*]  Ppuf_843-1200_[*bchX^Rs^*] with final 243 bp of *bchX^Rl^* fused at C-terminus cloned behind *bchYZ^Rl^* in pBBRBB[*bchYZ^Rl^*]  Ppuf_843-1200_[*bchX^Rl^*] lacking 243 bp C-terminus cloned behind *bchYZ^Rl^* in pBBRBB[*bchYZ^Rl^*] | This study  This study  This study  This study  This study |

**Supplementary Table 1. List of strains and plasmids described in this study**

†Department of Microbiology and Molecular Genetics, The University of Texas Medical School, Houston, Texas, USA

‡Department of Biochemistry, University of Oxford, Oxford, United Kingdom

| Primer name | Sequence (5’-3’) | Restriction site |
| --- | --- | --- |
| bciAKOCheckF | GATCTTCGTGCCCTTCGCGAAC |  |
| bciAKOCheckR | CATATCCTCGGCGATGCGCTC |  |
| bchYZKOCheckF | CTATGTGACGAACAACGTCTGC |  |
| bchYZKOCheckR | GTAGCACCTGCATATCCCATAACG |  |
| bchXKOUpF | CCGGAATTCCCGAAACTGATCGACAGTCTGG | EcoRI |
| bchXKOUpR | GATCGAGCCGCGCGAGGGTATCGATCATGCGGTGGCCCTCCAATCCAGGATC |  |
| bchXKODownF | GATCCTGGATTGGAGGGCCACCGCATGATCGATACCCTCGCGCGGCTCGATC |  |
| bchXKODownR | CCGTAAGCTTGGTGCAGCAGGCCGAACCCGAGATG | HindIII |
| bchXKOCheckF | CGCTCGATCTACGATGCGTCG |  |
| bchXKOCheckR | GTAGAAATGCGAGACGAACGAGAGG |  |
| bchNBKOUpF | GATCGAATTCGACACTTTCGGAATCGGAACG | EcoRI |
| bchNBKOUpR | CTGGACCTGAGACTTCACCGTGGCTCATTGCGCGGCCTCCATG |  |
| bchNBKODownF | CATGGAGGCCGCGCAATGAGCCACGGTGAAGTCTCAGGTCCAG |  |
| bchNBKODownR | GTCAAGCTTCAGTATTGCATGGACAGGAACC | HindIII |
| bchNBKOCheckF | CTCGCCGCAAAAGATGTAAGTC |  |
| bchNBKOCheckR | CATCTGCTCGAGGTTGTCGTC |  |
| bchCKOUpF | GACTCTAGATATCCGCTGATGGAACTCATG | XbaI |
| bchCKOUpR | TTCAGTTCGGGTGCATCCGTCATAGCGTTTTCCCCCGCGCTCT |  |
| bchCKODownF | AGAGCGCGGGGGAAAACGCTATGACGGATGCACCCGAACTGAA |  |
| bchCKODownR | GCCAAGCTTTCGTTCGATCCGACGAGGATCACCTTC | HindIII |
| bchCKOCheckF | TCAGTTCGATCTGGCGGTTC |  |
| bchCKOCheckR | CGATCTGATAGTTCGCCGACT |  |
| bchGSwapUpF | CATGGAATTCGAGACGCAGGTCTCGACGCTG | EcoRI |
| bchGSwapUpR | CCTGCAGATTGCTATTCATGTGTGTCACCTAAGCATGACG |  |
| bchGSwapDownF | CCTGGGCGATGCTGGTGGCAGCGATCGGACTCGGGTGACGCTGGGCTGGCTCG |  |
| bchGSwapDownR | GTCAAGCTTGCGAGGATCGTGAGGACAAGG | HindIII |
| bchGSwapbchG*^Rl^*F | CGTCATGCTTAGGTGACACACATGAATAGCAATCTGCAGG |  |
| bchGSwapbchG*^Rl^*R | TCACCCGAGTCCGATCGCTGCCACCAGCATCGCCCAGG |  |
| bchGSwapCheckF | GTCCGGAGTCGGGCATGCTG |  |
| bchGSwapCheckR | GACCTCGCGCAGCCCCTGAAGCAG |  |
| pK18*mobsacB*SequenceF | GAGTTAGCTCACTCATTAGG |  |
| pK18*mobsacB*SequenceR | CGGGCCTCTTCGCTATTACG |  |
| pK18*mobsacB*IntegrateF | GAGCGAAATGACCGACCAAG |  |
| pK18*mobsacB*IntegrateR | CTGCTTCGGGGTCATTATAG |  |
| pBBRBBSequenceF | GGTTGCGACACGCCATTCGTCG |  |
| pBBRBBSequenceR | GAGCCGTGTTTTCTGGACGATGG |  |
| bchYZ*^Rl^*F | GACAGATCTATGGCCAGCCTGACGGTGCC | BglII |
| bchYZ*^Rl^*R | GACACTAGTTCAGCGCTGCGCCGCCGTGG | SpeI |
| bchX*^Rl^*F | CGGCGCAGCGCTGAACTAGATTCGGCGAGAGGAAGGGAGA |  |
| bchX*^Rl^*R | GACTGCAGGTCCTGAAGTTAACTAGTTCATTCATAGAAGG |  |
| bchF*^Rl^*F | GCAGATCTATGAATTATACCCCCGAGC | BglII |
| bchF*^Rl^*R | GTCACTAGTTCAGCGACCGCTGCTGGCC | SpeI |
| bchC*^Rl^*F | GCAGATCTATGAAAACGAAAGCGATCGTG | BglII |
| bchC*^Rl^*R | GTCACTAGTTCAATGACCGGGGGCTCCGGAC | SpeI |
| bchG*^Rl^*F | GCAGATCTATGAATAGCAATCTGCAGG | BglII |
| bchG*^Rl^*R | GTCACTAGTTCACCCGAGGCCGATGGCC | SpeI |
| bchP*^Rl^*F | GCAGATCTATGTCCCGTGTGCTTGTGG | BglII |
| bchP*^Rl^*R | GTCACTAGTTCACCCGGAGGACGCCGAG | SpeI |
| minusN bchXRs F | GATCAGATCTATGGTCATCGCGATCTACGGCAAG |  |
| minusN bchXRs R | GATCACTAGTTCAAGCATCGTCGTAGATCAC |  |
| plusN in bchXRl N F | GATCAGATCTATGACGGATGCACCCGAACTG | BglII |
| plusN in bchXRl N R | GCCAGCATCCGTGGGGTCATCTGCGTCTTCTTCTTCGGCT |  |
| plusN in bchXRl X F | AGCCGAAGAAGAAGACGCAGATGACCCCACGGATGCTGGC |  |
| plusN in bchXRl X R | GATCACTAGTTCATTCATAGAAGGGCGCGG | SpeI |
| plusC in bchXRs X R | GAGGACTTTCGAATCCTGCGAAGCATCGTCGTAGATCACTTC |  |
| plusC in bchXRs C F | GAAGTGATCTACGACGATGCTTCGCAGGATTCGAAAGTCCTC |  |
| minusC bchXRl F | GATCAGATCTATGACCCCACGGATGCTGGC | BglII |
| minusC bchXRl R | GATCACTAGTTCAATAATCCGGGGAATTCAGATC | SpeI |

**Supplementary Table 2. List of primers used in this study**

Restriction enzyme cleavage sites used for cloning are highlighted in the primer sequence.

*bchY-bchZ*

ATGGCCAGCCTGACGGTGCCCGAGGGCAATCTGATCACCCCCGAGATCATCGAAACGCGCTCGAAATCCAGCGTCTCCGATCTCCAGAGCCAGACGGGCCATTCGGTCTCCCCGGATCATAACCTGACCCCGCAGACGATGTGCCCCGCCTTCGGGAGCCTCCGCGTCCTCACGCGGATCGATGGCGTCCAGACGGCCATGGTGACCGACACGGGGTGCCTGTACGGGCTGACGTTCGTCACGCATTTCTACGCGGCCCGCAAGAGCATCGCCGCCCCGGCCTTCGGGACCAAAGAACTCGTGGCGGGGAATGTCGCGGAAGCGGCGATCGCCGCGTGCATCGAAGCGAGCAAAATGCCGGGCACCAAACTGGTCGCGGCCATCTCGCTGTGCGTCGCGGAAACGGCCGGCCTGACCGAGGAAATGCTCCCGCGGCAGATCAACGGCGTGGATGTCATCCTCGTCCGGGTGCCGGCGTACGCGATCCATAGCCATCCCGAGGCGAAGGATGTGGCCATCAACGCCCTCCTCAAACGGTACGCGCCCGCCCAGACCGAAACCACCGAAGACGGGGTGGAGCGCCCCCTCACGTACGACGGGTCGAATGAGGAGGAGGAGGCCCTCCGCCGCAAGGT*CATCATGCTGGGGGAAGTCTTC*CCCGCCGATCCGATCTCCGTCGATGCGGTCCTCAAACGGCTGGGGGGCGGCCTGGCCGCGACCCTCCCCTCCCGGAATCTGGATGATTATAAACTCGCCCGCCGCGTCGGCGCGATCGCGGCCGTCCATCCCTTCTACGGGGGCTCGGTCGCCACCCTCCGCGCCGCCGGGGTCCCGATCATCTCCGGCGCGCCCGTCGGCCCGGAGGCCACGTACAACTGGATCAAGGCCGTCGCCGCCGCCCTGAAACTGGATGGCGAGCTGGCGGAGCAGGTGGCGGCCGAGGAGCGCGATAAGTGCAAAGCCATCATCTCCCAGTTCCCGCTGCGGGGGCGCGTGATGGTCTGCGGCTACGAGGGCAATGAGCTCCTCTATGCCCGGCTCCTGGTGGAGGCGGGGGCCGAAGTGCCGTATATCAGCACCTCGATCCAGAAATCCCTCTATACCGCGGCGGACGAGGCGTGGCTCAAAGCCCGCGGGACCCGGGAAATCGTCTATCAGAAGACCCTCGAGCAGGATGTCCGCGCGCTCGACCGCTATCAGCCCGACCTCGTCCTCGGCACGACCCCCCTGTCCGCGGAAGCCAAGCAGCGGGGGATCCCGGGCATGTA*CTACACCAATATGCTGTCGGTC*CGGCCCCTCTTCCTCTCGGCGGGCCTCGCGGGGACCATCCAGCTCATCCGGGACGCGCTGGAGCGCGCGCCGCGGTACGCCTGGATGCGCGAGTTCTTCGAGGGGCCGCGGCCCACGGCCCCGGCCGGCCCCGCGCCGATCAATACGATGGATTGACCGCGCCCCAGCGGCGTATAGACGGCACCGGCAATGACCGACCTCAACGATCTCCCGAACCGGGCCCCCGCGGCCGATCAGAGCCGGGCCCCCATCGAGCTCATCCGCGACCTGGAATCCACCAGCGGCTACTGGGCCGCGGTCTGGACCATGTGCACCCTGCCCGACGTGCACCTGATCTGCGATGCGCCCATCGGGTGCTTCAACCTCGTCGCCACGGCCGTGCCGGACTATACGGATGCCATCCCCCATATCCATAATATCACGCCCTCCGTCATGCGGGAGCAGGAAGTCACCCTCCTCGGCACGGCGGGCGCGGTCCGCCGCGCGGTCGAAGCCCTGCGGTCCCTCCACCCGGATAAAACGCTCATCGTCGTCTCGACGGCCGAATCGGAGATGATCTCCAGCGACCATTCGGATTGGCTCAGCAAGCTGGACCCCCCCGTGCCGTTCTTCTGGAGCCAGTCGCTCGAAGGCGACGAGTGGGAAGGGCGGGATCGCGCCCTCCTGTGGCTGTGGAACCGGTTCGGCCGCGAGCGGGCCGCCCCCCTCTCGGCCAATGATGCGCAGATGGTCAATATCATCGGCCCCACGTATGGCTGCTTCAACTCCCCCGCCGATCTCCATGAGATCAAGCGGCTGATCGAAGGCGCCGGGGGCAAGGTCAATCTCGTCTACCCGTTCGAAGCCACGCTCGAAGATACCCCCCGCCTCTCGCGGGGGGCGGTGAACGTGGTCATGTATCGCGAATTCGGGGAGGCGCTGGCCAAAGAACTGGGGCGCCCCTATCTGTTCGCGCCCATGGGCATCCGGGAGACCACGGAGTTCATCCAGCAGCTGGGCGACCTCCTCGGCACCCGGGCGCAGGCCGATGCGTTCATCGCCCGGGAGAAACAGACCACGCTGGCGCCCCTGTGGGATCTGTGGCGCGGGCCGCAGGGCGACTGGTTCGCCACCACCGAGTTCGCGGTCGTCGCGGGCCGGTCGTATACCGAAGGGCTCGTGCGCCTCCTGGCGGACGAACTGGGGATGAAACTCCATTTCGCCAGCGGCCGCCCCCGCCGCCCGGGGGAAATGGACAATATCCAGATCCGCGAAACGCTGCACAAGAAACAGCCGGCGTTCCTGTTCGGCAGCCTGAACGAAAAAATCTATCTGACGGAAGCGCAGGCGAAGGCGACGCATTTCATCCCGGCCGCGTTCCCGGGGGCGGTCGTGCGGCGGGCCCTCGGCACCCCGTTCATGGGGTATTCGGGCGTCGTCTATATCGTGCAGGAAATGGTCAACCGCTACTATGACCTCGTCTTCAATTTCCTGCCGTTCGACAACGTGGCCGCCGCCGGGCAGCCCAAGGAAGCGGCCAATCCGCTCACCGCGGCGAGCCGCCTCGTGTGGAGCGAAGCCGCCCGCGCGCAGCTGGATCGGCATCTCGAGGGCATCCCGTGGATCAGCCGCATCTCCGCGTCGCGGGAGCTCCGGGCCCAGGTCGAGGTGTATGCGATCAAACACAACCTCCGCGAAGTGACGCCCGACACGGTGGAGCAGGCCCTGGCCGCCACGGCGGCGCAGCGCTGA

*bchX*

ATGACCCCGCGCATGCTGGCGATCTATGGGAAAGGCGGCATCGGCAAATCCTTCTTCACCGCCAATCTGTCGGCCAAAATGGCCCTCAAAGGCCTGCGCGTCCTGCAGCTGGGCTGTGATCCGAAGCATGATTCGTGTAATGCCCTCTTCGAGGGCCGCAGCCTGCCCACCATCGGGGACCAGTGGCGCCTGTTCAAAGATGCCGGCCGCGAGAAAGAACTGGATGTCCGGCATCTGATCTTCAAAGCGGATCTCGGCCGCGGGGCGCAGCTGTTCGGCGCGGAACTCGGCGGGCCGGAAGTCGGGCGGGGGTGTGGCGGGCGGGGGATCTCGTTCGGGTTCGGGCTCCTGGAAGAACGCGGCATGGCGAATTGGGCGCTGGATTATATCATCATGGATTTCCTGGGGGATGTCGTCTGTGGCGGCTTCGCCACCCCGATCGCCAAATCGCTCGCCGAGGAAATCATCATCGTGGCGTCCCATGATCGGATGTCCCTGTATGCGGCGAATAACATCGCCCGGGCCGTGAATTATTTCCAGGCCATGGGGGGCTCCACGCGGCTCATCGGGATGGTGCTGAATCGCGATGACGGCAGCGGCGTGGCCGAACGCTTCGCGGAACGCATCGGGCTCCCCATCCTGGCGAAAATCCCCTTCTCGCCCGCGGCGCGGGCGCTCTCCGATCGCTGTAAACTCCTCTTCGAACTGCCGGAGATGGATGCGATCTTCGATGCCTTCGTGAATAAGATCCATCGGCGCGAATATACCGCGCCCACCAAAATCACCCCGCTCGAGTATGATGAGTTCCTCGCGGTGTTCGGGGCCAGCGAACCGCCCGATCTCCCCCAGGGCGCGACGCTCGATGAACTCATGGGGCGCGATAGCCAGTCGGTGGTGGGCATCGATCTGAATTCCCCGGATTATTCGCAGGATTCGAAAGTCCTCGTCCGGCGGATCATGCAGGAAATGGGGCTCAAAGTGACCCGCCTCGTGGAGGAACCGGAACGGGGGGTCGTGGTCACGACCGAAGCGGGGTGGGAGGCCGTCTTCGGCGATCCGTGTAAAGATATCGATGCCAAAATCGCCATCCTCTCGGCGCTCGGGCATTCCGGCGAGAAATTCCGCCTGGCCGATCTGCGGTATACGGCCGCGCCCTTCTATGAATGA

*bchF*

ATGAATTATACCCCCGAGCAGCTGGAACGGCGCAACCGCTCCTTCTGGACGCCCGTGCAGGCCGTCGGCGCGCTCGTGCAGTTCCTCACGTTCCTGGCGTCCCTGACCCTGGTCATCCGCTTCCTCACGACGGGCCAGGGGTATGAGGTCACGACGGTCGCCAATGTGGCGAAGGTGCTCATGCTCTATTTCATGACCGTCACCGGGATGGCGTGGGAGGATGAAGTGTTCGGCCGCATGTTCATGGCCAAAGAGTTCTTCTGGGAAGATGCGGGGAATCTGCTGTCGCTGGCGGGGAACACGGCCTACCTCGTCGCGCTCTTCGCCGGGGCCGACCATCGCACGCAGATGCTCGTCATGTGCGTCGCGCTCGCCACCTACGTCGTGAACTTCGTGCAGTTCGCCCGGCGCGGGGCCCAATCCGCGCGTCAAAAACGTGCCCGCGCCCTGGCCACGGCCAGCAGCGGTCGCTGA

*bchC*

ATGAAAACGAAAGCGATCGTGATCCCCGCCGCGCATACGGTGGAACTCCGGGAAGTCGAACTGAAACCGCTGGGCGCCGACGATGTCCTGGTGCAAACGACGCTCACGAGCATCAGCGCCGGGACCGAACGGATGCTCCTGCGCGGCGTCATGCCGCATCCCATGCTCCAGTTCCCCGTCGTCCCGGGGTATGAAACCGTCGGGCAGGTGATCGAAGCCGGCGCGAATGCCCGGGCGTGGCTCGGGAAACGGGTGTACGTGGGGGGGTCCTACGGCTTCGTCGGGGTCAATCCCGCCTTCGGGGGCCAGTCGGCGTACATCGTCACCCCCCAGTCGCACCTGACGGATCTGCAGAGCCTCACCGATGAACAAGGCGTGCTTCTCGCGCTTGCCGCGACGGCACTTCATGGTGTCGATGTGGCCTTTGGTGCGGGCCAAGCGTGTGGTGGAGAAGCTGCCCCTGAAGTTTGGGTGCTTGGTCAAGGCGTAGTGGGGCAACTTGCGGCGCGTTTTGCGAAAGCCCGTGGTGCGAGGGTCACGGTGTCGGATAAAGTCCCGTCGCGCCTCCGGTTGGCCGTCGCGGATGTGAAACTCCTGGCCGAGGACGGCGCGCTGAAAGCCGAAGATGGGTCCGCGGTGGCGGCCCTGGATAAAACGTGCGCGATCCTGATCGATGCGACGGGGAAGATGGAGGCCATCGCGCCGCGGCTCATGAATGTCCAGAAGGGCGGGCGGGTCGTGCTCCTCGGGTACTATGAACGGATCGACCTGCCGTATATGCCCGCGTTCCTGCGGGAACTGACCTTCGCCGTGAGCAAGGAATGGGCTCCTGGCGATCTTGCGAGAGCGCGTGATGCCATCGCCGCCGGCCAAGTGGAAGTGCGTTCCCTTATCACCCATCGGCTCCCGGCCGATGATGCGCCGCGGGCGTTCGATCTCGCCTTCAATGATCCGGACTGCGTGAAGATCACCCTCACGTGGTCCGGAGCCCCCGGTCATTA

bchG

ATGAATAGCAATCTGCAGGTCGCGGGCCACCCGCCGGCGAACAAACCGCTCCCCTTCTTCAAAGTGATGTTCACGCTGATGGAGCCGTACACGTGGTTCGCCCCCGCCTGGGCGTTCGTGGTCGGGTGCGTGGCCTCCCATTCCATCTATTTCGACCTCGGGGGGGATATCTGGCAGAGCGTCCTCAGCGTCGGGAAAATCGCCATCGGGACCCTCATGGCGGGCCCCCTGCTGGTCGGCTTCTCCCAGGTGTGGAATGATTGGTGCGACCGGGATGTGGATGCGATCAACCAGCCGGAGCGCCTGATCCCGAGCGGCCAGGCCACCCGGAATCAGGTCTTCGCGATCATCGCCATCCTGGGGGCGGCCGCCATGGCGATCGCGCTGTTCCTCGGGCCCCCGGTCGCCCTGGTCGCGCTGCTGGGGATCCTCATCGCGCTCGCGTACTCCGCGGAACCCATCCGGCTCAAAAAGAATGGGTGGCTCGGCAATCTCGCGATCGGCCTCACCTACGAGACGTGCGCGTGGATCGCGGGGCATTTCACCTTCGACCCCCAGCTGCAGTCCGCCGGCGCCTCCCGGTCGATGATCCTGGCCCTCATCTATGGCCTCGGCGTCACGGGGACCATGATCATCAACGACTTCAAATCGGTCGCCGGGGATCGCCAGATGGGGATCCGCTCGATCCCCGCGATGTATGGCGAATCGACCGCGGCCAAAATCGGCGTGGTCATCATCAATGTCGCCCAGCTGGTCGCGGTGGGCCTCCAGTTCGCGTGGGGGAATACGCTCGCCGCCGGGGCCTGCGTCCTGCTGCTCCTCGCGCAGCTGCCCATCCAGATGCAGCTCGTGCGCGAGCCCACCCAGCCGATGGCGGTGCGCTACAATATCGTGGGCATCCCGCCGTATACCTGGGCGATGCTGGTGGCGGCCATCGGCCTCGGGTGA

*bchP*

ATGTCCCGTGTGCTTGTGGTTGGTAGCTCGGTGGGGGGTTCGACCGCCGCCAATACGCTTCGGGACCTCGGGGTCGAGACGATCCTCCTCGAGCGGGACCTGAATTACGTGAAACCTTGTGGTGGGGCCGTGCCTCCCCGCGTGTTCAGCGACTGGGATCTGCCCCACGATCTGATCGATCGCAAAGTGACCATCGCCGTCGTGTGCTCCCCGCGGCATTACAGCGAATTCCCCGTGCTCTCCTCCCACCCGGCCCCCGAAACCGACTTCATCGCCATGGTCCGCCGCGAAAAATTTGATCGCTACATCCGGGATCGGGCGGTCCGGAAAGGGGCCGAACTGATCGAAGGCAAACTCGAGCATGTCGAATTCGGCCCCCGCGGCATCCGCGCCACCTACGAAGATAAACGGAACAAAACGCATATCATCGAAGCCGATGCGGTGATCGGGGCGGATGGGGCCTACTCCACCGTCGCGAAAAGCCTGGGCCTGGAGCGGCTCCCCATGGCGGTCGCGTACCAGGAGCGCATCCGGCTGCCCGCCCATGCGATGACGTATTGGGAAAAACGGGCGGCGCTCTACCTGGGGGACGATGTGTCCCCCGATCTCTATGCCTGGGTCTTCCCGAAATACGATCACGTCGCCGCGGGGATCGGGGCGGGCGCCGGGAAAACCAAAGCGGCGCGGCAGCTGCTCGCCAATCTCAAACATAAACTCCGGGATCAGCTGGGCGAAGGGCTGTCCGTCAAATTCGAAGCGCATCATCTCCCCATGCATCCGCGCAAACATCTCAGCTATGATCGTGCCGCGCTTGTCGGTGATGCCGCCGGTCTTGTGCAACAAACCTCCGGCGAAGGGATTTATTGGGCCATGAAATCCGGCGAAATGGCCGCGCGGGCGATCGCGAAACATCTGGATGCCCCGACCGCCGCGAATCTGCGGCAGGACTACGATGCGCCGTGGTGGAAAGCCTATCGCCCCACCTATCTCTTCCTGGCGTTTCTCCAGCGCATCTCCTACAACGGCGACCTCCAGCGCGAAATCTTCGCCAAAATGTGCGATGACCCCGATGTCCAGCGCCTCACGTTCGACGGCTATCTCACCAAGCACATCGAGCCCGCCCCCTGGACCGTGCAGATGCGCATCACCCGGCACTGGCTGTCGACGGCGGTGCATGAATGGATGCGGCAGCAGAAAGAGAAGACCTCCTCGAGCTCGGCGTCCTCCGGGTGA

**Table 3. “*Ca*. Roseilinea sp. NK_OTU-006” genes codon-optimised for *Rba. sphaeroides***
